# pH-driven dynamics and mechanisms of LNP-mediated siRNA delivery

**DOI:** 10.64898/2026.02.11.705380

**Authors:** Kazi A. Hossain, Mariana Valério, Pedro Medina, Paulo C. T. Souza, Modesto Orozco

**Affiliations:** Institute for Research in Biomedicine (IRB Barcelona), The Barcelona Institute of Science and Technology (BIST), 08028 Barcelona, Spain; Laboratoire de Biologie et Modélisation de la Cellule, CNRS, UMR 5239, Inserm, U1293, Université Claude Bernard Lyon 1, École Normale Supérieure de Lyon, 46 allée d’Italie, 69364, Lyon, France; Centre Blaise Pascal de Simulation et de Modélisation Numérique, École Normale Supérieure de Lyon, 47 allée d’Italie, 69364, Lyon, France; Department of Biochemistry and Biomedicine, Faculty of Biology, University of Barcelona, 08028 Barcelona, Spain

## Abstract

Lipid nanoparticles (LNPs) are a transformative technology for RNA delivery. Here we present a comprehensive analysis of a clinically relevant DLin-MC3-DMA (MC3)-based LNP formulation that exhibits strong short interfering RNA (siRNA, ∼21 nt) delivery and potent inhibition of targeted RNAs. Using a combination of coarse-grained and atomistic molecular dynamics simulations, we characterize the pH-dependent structure of both unloaded and siRNA-loaded LNPs, elucidate the mechanism of siRNA encapsulation, and use these models to propose a plausible mechanism of endosomal escape. Consistent with prior experimental and computational studies, our simulations reproduce an inverted-hexagonal– type morphology in a ∼70 nm siRNA-loaded LNPs. For the smaller LNPs mimetics, with only one siRNA sequence, we observe a similar hydrated core as observed in the larger 70 nm LNP model. Such hydrated channel provides a polar environment suitable for accommodating RNA, whose charge is neutralized mainly by the MC3 ionizable lipids that remain protonated near the RNA even at high pH, thereby bridging the RNA with the surrounding lipid environment. This structural picture is also consistently observed across our multiscale simulations, with smaller self-assembled LNP mimetics reproducing the same local organization at both coarse-grained and atomistic resolution. A potential mechanism of endosomal escape, emerges spontaneously from the simulations, involving stalk formation between the LNP and the endosomal membrane, followed by the opening of a water-filled pore that permits slow siRNA diffusion, in line with the low efficiency and slow kinetics reported for endosomal escape. The rate-limiting step of endosomal escape arises from persistent electrostatic coupling between RNA and protonated MC3 ionizable lipids maintained by the immediate RNA–lipid environment, hindering cargo disengagement even after pore formation. This delayed release is consistent with the experimentally observed time-dependent inhibitory activity of the loaded LNP. Together, these results highlight the importance of local pKₐ and protonation effects near the RNA, which are not captured by global apparent pKₐ measurements.

## Introduction

The emergence of nucleic acid (NA)–based therapeutics, including antisense oligonucleotides, aptamers, microRNAs, messenger RNAs (mRNAs), RNA sponges, and RNA interference (RNAi) strategies, has revolutionized the treatment landscape for a wide range of diseases, from infectious and neoplastic disorders to rare genetic conditions^1–5^. Unfortunately, despite their therapeutic potential, the clinical translation of these molecules remains limited by formidable delivery challenges. Unmodified, naked NAs are inherently unstable in biological environments, prone to rapid enzymatic degradation, exhibit poor cellular uptake, and are rapidly cleared from circulation^6^. Early delivery approaches, such as chemical modification or conjugation with peptides, sugars, or lipids, provided only partial solutions, offering modest protection and uptake, while often introducing toxicity or interfering with molecular mechanisms involved in NA therapeutic activity^7,8^. These limitations have underscored the need for delivery systems that can simultaneously protect RNA, facilitate intracellular transport, and remain biocompatible and scalable for clinical use. In this context, lipid nanoparticles (LNPs) have emerged as a transformative technology for RNA delivery^9–11^. LNPs offer high encapsulation efficiency, favorable biocompatibility, and improved endosomal escape kinetics, making them the leading platform for the clinical application of RNA-based therapeutics^12,13^.

LNPs are typically composed of four key components—an ionizable lipid, sterol (e.g. cholesterol - CHOL), a helper phospholipid, and a polymer-conjugated stealth lipid (e.g., PEGylated lipid)—each fulfilling a distinct structural and functional role. Ionizable lipids such as DLin-MC3-DMA (MC3), ALC-0315, and SM-102 enable efficient RNA encapsulation through electrostatic complexation at acidic pH, while minimizing systemic toxicity due to charge neutralization at physiological pH. Helper phospholipids and sterols stabilize the nanoparticle structure and modulate membrane fluidity and internal organization, whereas PEG-lipids control particle size, colloidal stability, and *in vivo* circulation time^11,14,15^. However, despite their remarkable success, fundamental questions about the behavior of LNPs as vehicles for NA delivery remain unresolved, including their internal architecture, pH-dependent structural reorganization, and the molecular basis of endosomal escape. This limited mechanistic understanding hampers the rational design and optimization of next-generation LNPs, leaving the field reliant on high-throughput combinatorial screening and empirical discovery^16,17^. While efforts to apply artificial intelligence (AI) to LNP development are promising, they are constrained by the scarcity and heterogeneity of annotated experimental datasets, and they rarely provide insight into underlying molecular events such as local protonation, transient fusion, or phase transitions^18–22^.

Current knowledge of LNP structure and behavior has been primarily derived from experimental methods such as small-angle X-ray scattering (SAXS), cryo-electron microscopy/tomography (cryo-EM/cryo-ET), and packing-sensitive spectroscopic probes^23,24^. These studies have revealed that LNPs can adopt diverse internal morphologies—including lamellar, dense-core, inverse hexagonal, or worm-like inverse phases—whose prevalence depends strongly on formulation composition, pH, and cargo^25–27^. Complementary cell biology experiments employing fluorescent probes or reporter-gene assays with encapsulated nucleic acids have further provided valuable insights into LNP uptake and endosomal escape, including comparisons between standard and peptide-decorated LNPs^28–33^. However, experimental approaches alone cannot capture the vast compositional landscape of possible lipid formulations or the dynamic structural and compositional changes that occur during intracellular trafficking. Molecular simulations offer a powerful complementary strategy to bridge these gaps: coarse-grained (CG) molecular dynamics (MD) simulations can probe realistic LNP size and assembly dynamics, while atomistic (AA) MD simulations enable detailed investigation of local packing, interfacial organization, and pH-dependent structural transitions^34–37^. Nevertheless, most published simulation studies to date rely on oversimplified or bilayer-mimic LNP models^35,38–44^, often neglecting explicit protonation equilibria and providing limited mechanistic insight into critical dynamic processes—such as structural reorganization, membrane fusion, and RNA release from endosomes—a key yet poorly understood step that ultimately determines whether the therapeutic oligonucleotide reaches the cytosol or is degraded within lysosomes^15,45–49^.

We present here a multiscale study investigating the coupling between protonation state, cargo, and LNP structure, with a particular focus on how these factors influence cellular trafficking. Thus, using CG MD simulations, we characterized the detailed architecture of a full-scale MC3 based-LNP model (approximately 70 nm in diameter) in aqueous solution, which we have experimentally demonstrated to show excellent encapsulation and delivery properties^50^. Furthermore, we construct a smaller system that reproduces key features of the internal organization as the full-scale LNP model, using it to analyze the influence of cargo on nanoparticle structure and the mechanism of pH-dependent siRNA encapsulation. Transitioning from CG to atomistic resolution, we employed constant-pH molecular dynamics (CpHMD) to characterize, how the LNP structure varies along a broad range of pHs. Such an analysis is particularly relevant for MC3-like LNPs because the protonation state of the ionizable lipid is highly sensitive to its local environment. While the intrinsic aqueous pKₐ of MC3 has been reported to be approximately 9.47^51^, experimental measurements yield an apparent pKₐ of about 6.44^11^ in LNP formulations, reflecting the strong influence of the environment on lipid protonation. These observations suggest that protonation behavior may vary significantly across different regions of the particle. CpHMD simulations therefore provide a specific opportunity to directly investigate how local environments within the LNP modulate protonation states, enabling, the determination of the (apparent) pKa distribution within the LNP. By combining CG and atomistic simulations, one plausible sequence of molecular events associated with the endosomal escape of siRNA in this MC3–based LNP system is found. Accordingly, the LNP first forms a shallow fusion stalk with the endosomal membrane, followed by the transient opening of a water-filled channel that can enable siRNA to move toward the cytosol. The simulations further suggest that persistent interactions between siRNA and protonated ionizable lipids may also contribute to the endosomal escape bottleneck observed in MC3-based siRNA LNPs. These results highlight the power of simulations to characterize the basic mechanisms of LNP encapsulation and delivery and support the use of simulations to help in the design of new LNP with similar formulation and cargo.

## Results and discussion

### Palmitate-containing LNPs encapsulate and deliver siRNAs with high efficiency

We recently developed a distinct LNP formulation for the delivery of therapeutic siRNAs to HER2⁺ breast cancer cells^50^. The optimized formulation included: (i) the neutral lipid DSPC (1,2-distearoyl-sn-glycero-3-phosphocholine), (ii) cholesterol, (iii) the ionizable lipid DLin-MC3-DMA, and (iv) the PEGylated lipid DPG-PEG (1,2-dipalmitoyl-rac-glycero-3-methyl poly-oxyethylene-2000), combined at a molar ratio of 10:38.5:50:1.5 (DSPC:CHOL:MC3:DPG-PEG). In contrast to the canonical DLin-MC3-DMA LNP formulation, which typically employs DMG-PEG, the formulation used in this study contains DPG-PEG. This composition was chosen to match our previously developed and experimentally validated LNP formulation^50^. HER2-silencing siRNA was incubated with the lipid mixture at pH 4, using a siRNA:lipids mass ratio of 1:16.7, corresponding to a N/P (nitrogen/Phosphorus) ratio of ∼5. This process resulted in efficient siRNA loading, with encapsulation efficiency exceeding 80% (Fig. 1a). Following extrusion, the resulting LNPs displayed a uniform hydrodynamic diameter of approximately 76 nm (Fig. 1b). These siRNA-loaded LNPs remained stable at physiological pH and successfully crossed the plasma membrane of HER2⁺ breast cancer cells, enabling efficient intracellular siRNA delivery (<u>Fig. S1</u>) and subsequent knockdown of the targeted genes^50^. Thus, this LNP formulation represents a robust and highly efficient delivery platform and will serve as the model system for all simulations described below.

**Fig. 1:**
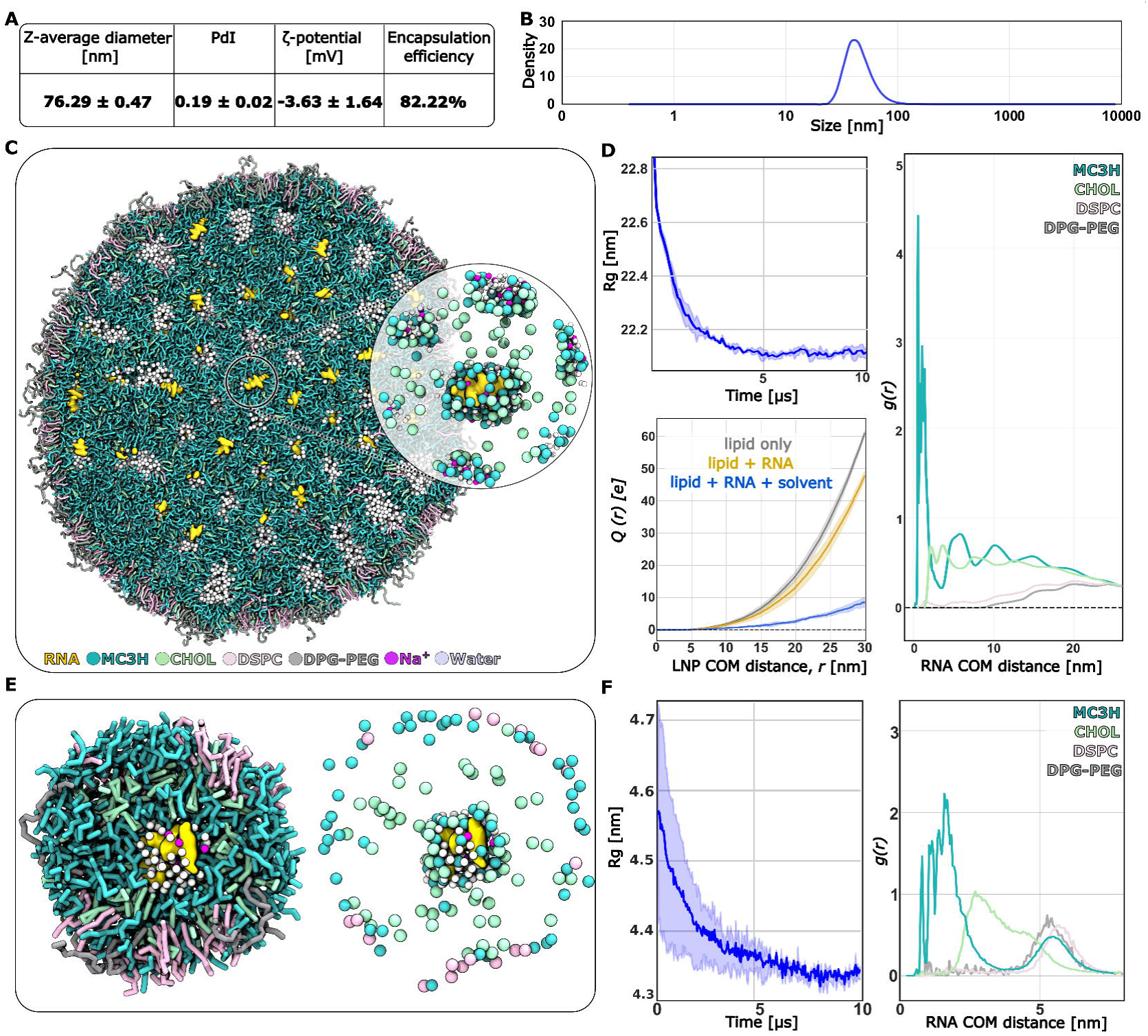
Structural validation and global organization of the large and small HER2–siRNA LNP models. (A) Summary table of experimental characterization: z-average diameter, polydispersity index (PdI), measured ζ-potential and siRNA encapsulation efficiency. Note that the experimental hydrodynamic radii corresponds to the PEG-coated particle plus the solvation shells. (B) Intensity-weighted hydrodynamic size distribution measured by DLS for the experimental formulation (DSPC:CHOL:MC3:DPG-PEG = 10:38.5:50:1.5). (C) Coarse-grained representative structure of the large HER2–siRNA LNP after multi-µs sampling, showing RNA (gold), protonated MC3H (cyan), CHOL (mint), DSPC (pink), DPG-PEG (gray), Na⁺ (magenta) and water (pale blue). The inset highlights the hydrated, MC3-stabilized core and nearby counterions around a RNA. (D, left) Time series of the radius of gyration (Rg) for the large LNP showing rapid compaction and convergence during the coarse-grained run. Data presented as mean ± SD over three replicas shaded, see Methods. (D, bottom-left) Radial cumulative charge profiles Q(r) computed from three selections: lipids only (grey), lipids + RNA (gold), and lipids + RNA + solvent/ions (blue). Including solvent ions strongly reduces the net enclosed charge near the particle, consistent with a near-neutral electrokinetic signature from our experimental results. Data presented as mean ± SD over three replicas shaded. (D, right) Radial distribution functions g(r) of lipid headgroups around the RNA center-of-mass (COM): MC3H is strongly enriched at short distances (core-proximal), CHOL shows enhanced density at intermediate distances, whereas DSPC and PEG-lipids populate the outer shell. (E) Left: representative coarse-grained structure of the smaller, self-assembled LNP (∼10 nm diameter); right: same small particle shown as a bead representation highlighting water and ion distribution around the core. (F) Left: Rg time series for the small self-assembled particle. Data presented as mean ± SD over three replicas shaded; Right: g(r) of lipid headgroups around the RNA COM for the small particle — the same spatial ordering (MC3H core, cholesterol/intermediate, DSPC and PEG outer shell) is preserved.

### Validated large-scale model reveals stable inverted-hexagonal morphology of HER2– siRNA LNP

To gain structural insight on our LNP, we first constructed a realistic-size prebuilt hexagonal model (see Methods) of a HER2-silencing siRNA encapsulated within a lipid formulation consistent with our experimental composition (see above). The intrinsic pKa of MC3 is 9.47^51^, and since LNP assembly occurs (see above) under acidic pH (∼4) we consider all MC3 molecules in their protonated, cationic form (hereafter referred to as MC3H). The total system contains 422,800 beads, corresponding to on the order of ∼4 million atoms (including hydrogens) at atomic resolution. Then, we carried out multi-microsecond coarse-grained molecular dynamics (CG-MD) simulations (see Methods), allowing the system to relax spontaneously so that its final structure was determined by the underlying physics. Over the course of simulations, we observed that in around 5 µs (CG-time) the particle rapidly compacted into a roughly spherical structure with an overall maximum diameter of ∼72 nm (very close to the experimental hydrodynamic diameter; <u>Fig. S2</u>a), while maintaining an inverse hexagonal phase (H_II_)-like internal organization (Fig. 1c, <u>Fig. S2</u>b,c and also see <u>Movie</u> <u>SM1</u>), which has also been proposed experimentally for MC3-based LNPs at acidic pH values^26,52,53^. The evolution of the radius of gyration (Rg) (Fig. 1d, top-left) confirmed structural convergence toward a stable, compact particle, not far from the experimental estimates (see Fig. 1a,b). These simulations were performed in triplicate, with all simulations consistently converging to the same structures. The structural organization of the system was further characterized by calculating the SAXS profile after 10 μs of simulation (<u>Fig.</u> S2D). The profile exhibits four reflections at q = 0.203, 0.320, 0.438, and 0.555 nm^-1^, corresponding to relative peak positions of 1.00, 1.58, 2.16, and 2.74, respectively. These peak positions show approximate agreement with the characteristic reflections expected for a two-dimensional hexagonal lattice, whose ideal relative positions are 1.00, 1.73, 2.00, and 2.65. The calculated profile is therefore consistent with hexagonal ordering and agrees with experimental observations for similar MC3-based compositions, supporting an HII-like internal organization.

To understand in detail the internal organization of the LNP, we computed radial distribution functions (RDFs) of the headgroup beads (see Methods) of individual lipid species as a function of distance from each RNA molecule and then averaged over all RNAs and trajectories. Results in Fig. 1d, right show that the positively charged MC3H lipids were preferentially localized near the RNA (within a 3 nm radius), forming direct and water-mediated contacts that effectively neutralize the negative RNA phosphate backbone. A flatter MC3H distribution (Fig. 1d, right) at larger distances (beyond ∼20 nm radius) indicates that a fraction of protonated lipids resides near the surface, where they interact with water and help to stabilize the inverted-hexagonal morphology in the presence of RNA. Cholesterol (CHOL) exhibited enhanced density near RNA (1-5 nm) but displayed a broad distribution consistent with its dual role in packing between hydrophobic tails and stabilizing the particle core^54,55^. In line with previous simulations and experimental studies, PEG-lipids (DPG-PEG) remained predominantly at the particle surface (stay at >15 nm radius; Fig. 1d, right) and formed an extended PEG brush-like corona (Fig. 1c), providing colloidal stability and preventing aggregation^35,56,57^. The helper phospholipid DSPC was also enriched at the outer interface (Fig. 1d, right), forming a quasi-continuous shell between the aqueous phase and the hydrophobic interior, consistent also with earlier observations ^38^.

The water and ion density profiles (<u>Fig. S3</u>) further characterize the H_II_ morphology: the LNP core remained hydrated, providing a polar microenvironment suitable for accommodating RNA, while the surrounding shell excluded bulk water. To quantify electrostatic balance, we calculated the radial charge-density profiles (as a function of distance from LNP center) for (i) lipids only, (ii) lipids + RNA, and (iii) the full solvated system (<u>Fig. S4</u>). The lipid-only profile exhibited a quite high net positive charge density of ∼0.6 e nm^-3^; inclusion of RNA resulted in partially neutralizing this charge (∼0.4 e nm^-3^), and solvent counterions further screened the potential, yielding an overall near-neutral formulation. Thus, the calculation of the enclosed charge (Q(r)) of the loaded LNP as a function of distance from the LNP center (Fig. 1d, bottom-left) shows that the interior region is essentially charge-neutral, and only beyond ∼15 nm does the profile deviate slightly, rising to a small positive value of a few elementary charges. Using a simple spherical electrostatic estimate to convert Q(r) into an approximate potential, the corresponding ψ(r) at plausible slip-plane radii (8–12 nm) falls in the low-millivolt range—of the same order as our experimentally measured ζ-potential (Fig. 1a), showing that the overall electrostatic environment predicted by the CG model is compatible with the weakly negative ζ-potential we observe experimentally (Fig. 1a).

### Simplified self-assembled LNP preserves key structural features of the larger model

To probe pH-dependent structural reorganization and to bridge between atomistic resolution and mesoscale dynamics, we next developed a smaller model that can be treated at the atomistic level retaining the essential physicochemical characteristics of the experimentally validated large LNP. This model differs from experimentally tested LNPs in several respects, namely the reduced particle size, the siRNA copy number, and the absence of full mesoscale morphology characteristic of larger LNPs. Rather than reproducing the complete architecture of experimental LNPs, the purpose of this reduced system is to reproduce the local structure of lipids around RNA, providing atomistic insight into siRNA–lipid interactions, hydration effects, and pH-dependent structural reorganization within feasible computational timescales.

To ensure that this system would tend to form such organization spontaneously, the smaller LNP (∼10 nm in diameter) was generated by self-assembly from randomly distributed lipid and siRNA components in a simulation box (see Methods). The formulation and stoichiometry were identical to our experimental composition (DSPC:CHOL:MC3:DPG-PEG=10:38.5:50:1.5), but with the system containing only a single siRNA molecule (see Methods).

Within ∼7 μs of CG MD simulation, spontaneous organization of the components led to the formation of a compact lipid assembly around the RNA (<u>Movie SM2</u>). At this reduced scale, the global distinction between inverted-hexagonal and inverted-micellar morphologies becomes less pronounced; instead, the key conserved feature is the local RNA-centered packing motif. This local RNA-proximal organization closely resembles the RNA–lipid environment observed in the large LNP model (see Fig. 1c and 1e for comparison) despite differences in overall particle size and RNA content. The temporal evolution of the radius of gyration (Rg) (Fig. 1f left) confirmed the formation of a compact, stable particle. The resulting model effectively resembles the RNA local environment on the large LNP (Fig. 1c, *inset*), where the siRNA remains enclosed within a hydrated polar core, shielded by the MC3H lipids (Fig. 1f, right). A detailed structural comparison around a single siRNA molecule in the full-size and reduced LNP shows similar lipid organization (see <u>Fig. S5</u> and Fig. 1f, right), confirming that the smaller self-assembled particle preserves the same internal packing, charge screening (see <u>Fig. S6</u>), water and ion density profiles (see <u>Fig. S7</u>) as the realistic large-scale LNP. This structural consistency establishes the simplified LNP as a valid and efficient model for probing pH-dependent protonation equilibria, structural rearrangements, and membrane interactions discussed in the following sections.

To further verify that the assembly protocol does not introduce structural bias into our larger models, we constructed a prebuilt small LNP using the identical protocol applied to the full-sized system. Because this prebuilt model bypasses the initial self-assembly phase, it reaches equilibrium more rapidly; we therefore evaluated its structural organization in triplicate for 5µs. Comparative analysis of the internal organization and shape (Fig. S8), confirmed that the prebuilt small LNP converges to the same stable architecture as the spontaneously self-assembled system (Fig. 1F, left panel).

### Multiscale simulations sampling reveals pH-dependent organization and variable pKa in the ionizable lipids depending on RNA distance

To characterize the pH-dependent organization of the LNP we first backmapped (see Methods) the self-assembled CG model of the small LNP (Fig. 1e) to an atomistic representation and equilibrated it (<u>Movie SM3</u>). The resulting system contains around 414,908 atoms placed in a cubic box of approx. 16.1 × 16.1 × 16.1 nm^3^ of volume. We then performed atomistic CpHMD simulations by making all MC3 lipids (300 in total) titratable and running two replicas at each pH value from 4 to 14 (see Methods) until the protonation fractions reached convergence (see Methods and <u>Fig. S</u>9).

Representative structures at different pH values are shown in Fig. 2a, *top* (see also <u>Fig. S</u>10). At acidic pH, most MC3 molecules remain protonated (MC3H), and the particle adopts a less compact morphology that is slightly elongated in shape (Fig. 2a, *top*). As the pH increases, MC3 becomes progressively deprotonated, with a sharp transition occurring at pH 7.7 (Fig. 2b), not far from the apparent pKa (∼6.4) measured for a similar MC3-based LNP^11^. This transition (Fig. 2a, *top* and b) coincides with a marked structural change: the LNP becomes more compact and spherical; but irrespective of the pH, the core of the LNP remains always hydrated (<u>Fig. S</u>11). This trend is consistent with experimental observations from a cryo-EM study showing that LNPs are more hydrated and less compact at acidic pH and become more ordered and compact near neutral pH^26^.

**Fig. 2:**
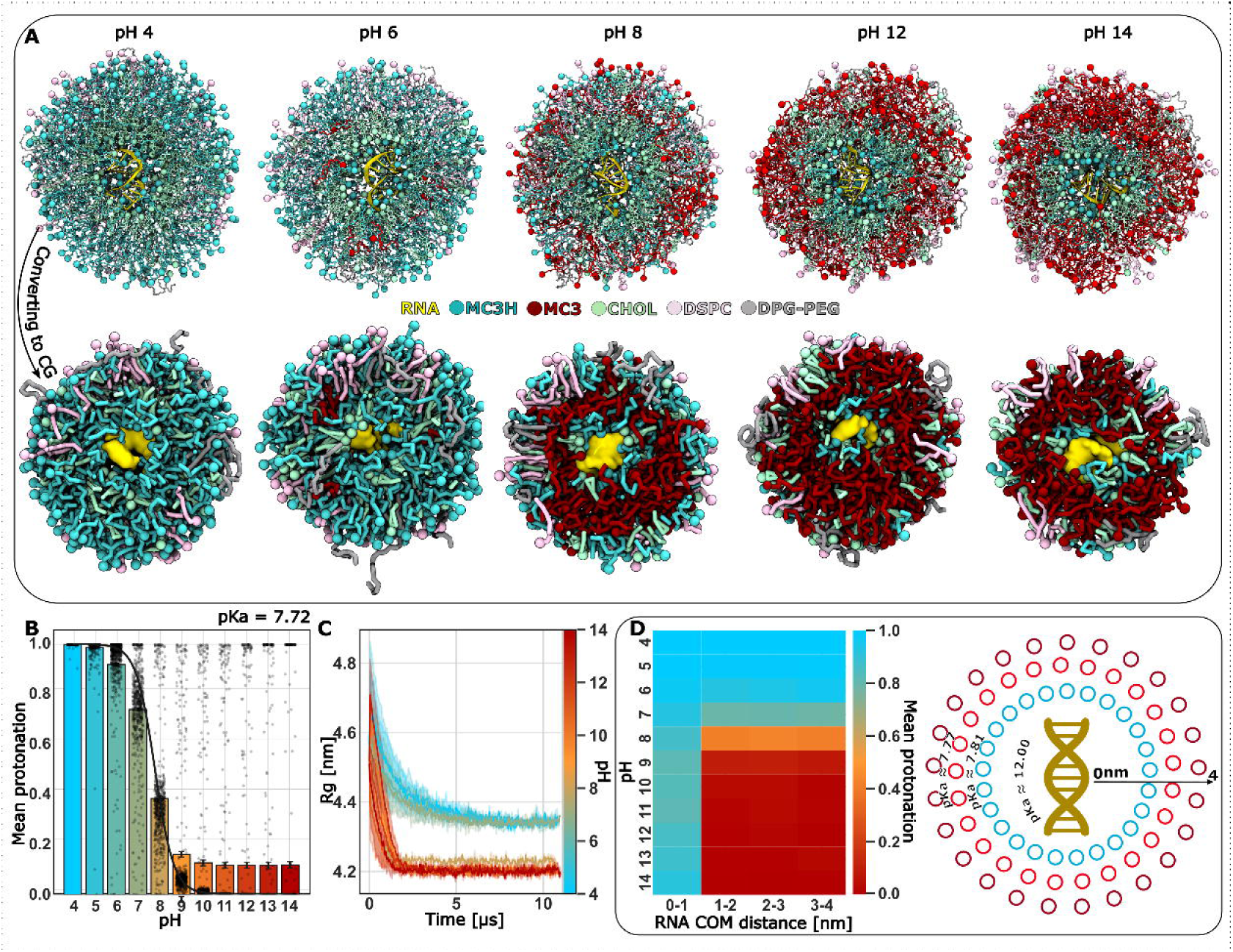
pH-dependent protonation and structural reorganization of the small HER2–siRNA LNP. (A) Representative atomistic (top row) and coarse-grained (bottom row) structures of the smaller self-assembled LNP at different pH values (acidic to alkaline), showing RNA (gold), protonated MC3H (cyan), neutral MC3 (red), CHOL (mint), DSPC (pink), DPG-PEG (gray). Increasing pH drives progressive MC3 deprotonation at the particle periphery and a concomitant compaction of the LNP (compare pH 4 with pH 8). The coarse-grained views (bottom) show the same qualitative reorganization while allowing longer sampling of mesoscopic rearrangements. (B) Mean protonation fraction of all MC3 residues as a function of pH. Data presented as bar plot of mean protonation ± SD (n=596). Fitting the titration curve with the Henderson–Hasselbalch equation gives an apparent global pKa ≈ 7.72 for MC3 in this siRNA-loaded formulation. The convergence of mean protonation and pKa is presented in <u>Fig. S</u>9. (C) Radius of gyration (Rg) time series from extended CG sampling; colored by pH scale. Higher pH produces systematically smaller Rg (more compact particle). Data presented as mean ± SD over two replicates. (D) Spatially resolved protonation. Left: heatmap of mean MC3 protonation fraction binned by radial distance from the RNA center-of-mass (COM) and by pH. Even at very high alkaline pH, MC3 residues within 1 nm of the RNA remain highly protonated. Right: schematic circle plot showing how the distance-dependent protonation to local apparent pKa values: the inner shell around the RNA has a locally raised pKa (∼12), whereas the outer shells approach the global pKa (∼7.7). This radial heterogeneity explains how a strongly protonated inner shell can stabilize RNA encapsulation while the particle surface becomes neutral at physiological pH. Colored circles (red, orange, and blue) indicate the mean protonation state at pHs between ∼8 and 9, with higher mean protonation closer to the RNA and decreasing protonation with increasing radial distance.

Although the atomistic CpHMD runs converged, the mean protonation fractions within a shorter ∼400 ns (<u>Fig. S</u>9), those trajectories might be short to represent collective lipid rearrangements. Thus, to explore potential protonation-dependent structural perturbations on longer timescales, we mapped (see Methods) the equilibrated atomistic models to CG representations (Fig. 2a, *top*) and extended sampling up to ∼10 μs (CG-simulation time; see Methods). Despite the resolution change and the extension of trajectories, the CG simulations reproduced the same qualitative trends observed in the atomistic snapshots: with increasing pH the particle becomes more compact, whereas acidic pH favors a looser, less compact morphology (Fig. 2a, bottom). The representative structure from CG (Fig. 2a, bottom) show larger amplitude rearrangements of MC3 residues than atomistic, but the pattern is the same — MC3 residues near the RNA remain cationic (MC3H), while peripheral MC3 gets deprotonated and migrates outward as pH rises. This behavior is reflected in the radius of gyration (Rg) time series (Fig. 2c): Rg values are systematically higher at low pH (less compact) and decrease toward higher pH (more compact), consistent with the atomistic trends described above.

Finally, to capture spatial heterogeneity of protonation within the LNP, we partitioned interior of the particle into concentric distance bins with respect to RNA center of mass (COM) and computed the mean protonation fraction of MC3 residues as a function of both pH and radial bin (Fig. 2d, left). Interestingly, MC3 residues within the innermost bin (0–1 nm from RNA) remain essentially fully protonated (MC3H) at all pH values sampled. In contrast, MC3 residues located farther from RNA showed pronounced pH dependence and lost protonation above pH 7 (Fig. 2d, left). These observations indicate that the RNA microenvironment strongly stabilizes the protonated form of the ionizable lipids (MC3H), whereas the outer layers are more prone to change in the protonation state. We then translated these distance-dependent protonation profiles into local apparent pKas by fitting the titration curves for each radial bin using the Henderson–Hasselbalch equation (Fig. 2d, right). This yielded a high apparent pKa (∼12) for MC3 residues in the vicinity of RNA and a substantially lower apparent pKa in the outer region, which approaches the global value of ∼7.7 obtained from the overall titration curve (Fig. 2b). This radial variation in pKa reflects the strong electrostatic stabilization provided by the RNA phosphates and the polar environment within the core (<u>Fig. S</u>12). Together, these results provide a unified explanation for two observations: (i) siRNA remains stably encapsulated at physiological or serum pH because the inner MC3H layer does not deprotonate even at high pH; and (ii) the LNP surface becomes neutral at serum pH, making the formulation safer by reducing unwanted interactions^51^.

### Cargo affects LNP morphology and protonation

To test the impact of siRNA cargo on LNP organization and protonation, we generated self-assembled “empty” LNP model (same lipid composition of the small LNPs as before, but no RNA) in multiple replicas and analyzed their structure and titration behavior (<u>Fig. S</u>13 and <u>Movie SM4</u>) in the same way as for the siRNA-loaded particles (see Methods). Structurally, the empty particles displayed a markedly different internal morphology: instead of forming one compact core as seen in the siRNA-loaded LNP, the empty LNPs show multiple internal water pockets, including one larger central pocket (<u>Fig.</u> <u>S</u>13). This morphology may represent an early stage reorganization toward an inverted-hexagonal-like structure, which could stabilize given the small LNP model. Nevertheless, it is consistent with some experimental observations of empty or cargo-deficient formulations^52,58,59^. This difference indicates that the presence of a polar, highly anionic cargo strongly biases the internal packing toward a pronounced RNA–MC3H–stabilized hydrated core (compare Fig. 1e and <u>Fig. S</u>13), with the nature of the resulting internal organization expected to depend on cargo size and charge density.

We next back-mapped representative empty-LNP CG structures to atomistic resolution and performed CpHMD simulations over the same pH range (4–14) as described above. Similar to the siRNA-loaded system, the empty LNPs showed increasing compaction with increasing pH; however, their internal organization remained qualitatively distinct (<u>Fig. S</u>14a). Water-density profiles support this trend, showing progressive dehydration of the particle with increasing pH, particularly beyond ∼4 nm from the center (<u>Fig. S</u>14b). From the global titration curves, we obtained an apparent pKa of 7.48 for MC3 residues in the empty formulation (<u>Fig. S</u>14c), which is close to the value of 7.72 for the siRNA-loaded particle (Fig. 2b). Thus, the presence of a single siRNA molecule does not substantially shift the global apparent pKa of MC3 in this lipid composition, but the protonation landscape differs significantly. Thus, distance-resolved protonation analysis shows that MC3 residues in the inner region of the empty particle remain protonated, but to a noticeably lower degree than in the siRNA-loaded core (compare first column of heatmap in <u>Fig. S</u>14d, left and Fig. 2d, left). In other words, while the empty LNP retains some protonation in its center, it does not develop the persistent, highly cationic MC3H shell that forms around RNA. Translating these profiles into local apparent pKa values further emphasizes this difference: although the very center retains a high pKa (∼11), (<u>Fig. S</u>14d, right), the pKa drops sharply just outside this region (<u>Fig. S</u>14d, right). The presence of polar patch or water accumulation in the center of the empty nanoparticle justifies this small spot of high apparent pKa in the absence of RNA (see <u>Fig. S</u>13).

Together, these results indicate two coupled cargo effects. First, the negatively charged RNA organizes nearby MC3H into a stable, highly protonated inner shell, which in turn stabilizes a hydrated core—an arrangement compared to the empty particles, but larger to adapt to the cargo size. Second, despite these local differences, the cargo may only modestly influence the global apparent pKa of MC3 residues, showing that global titration curves can mask local, functionally important protonation differences that are directly relevant for stability and pH-responsive behavior.

### LNP-endosome interactions induce stalk intermediates and precede pore opening

To understand the behavior of LNPs inside an endosome, and motivated by a plethora of experimental data showing PEG-lipids are not internalized into the endosome ^60–62^ we used our reduced model of LNP to create a naked (i.e PEG-free) LNP and placed it in the interior of a model endosome. Hence, the present model does not capture the complete intracellular pathway and is intended to represent a later stage of intracellular delivery, after protein corona interaction, PEG-lipid shedding, and endosomal maturation has occurred.

In particular, we selected the representative pH 6 structure (as the early endosomal pH ≈ 6.0)^63^ from our CpHMD calculations and embedded the resulting LNP inside a prebuilt endosomal vesicle (<u>Fig. S1</u>6) with a realistic heterogeneous lipid composition^64^, in which anionic lipids are primarily present in the luminal (inner) leaflet of the endosomal membrane, where they are thought to promote electrostatically driven membrane remodeling and fusion processes ^37^. The final systems were then subjected to CG MD simulations (see Methods), which show that the naked LNP rapidly reoriented inside the vesicle and moved toward the inner leaflet (<u>Movie</u> <u>SM5</u>). Within ∼1.3 µs (CG-time) the particles established extensive lipid–lipid contacts (“lipid handshake”), followed shortly by the formation of a shallow fusion stalk (Fig. 3, *top*). The stalk is the canonical first intermediate of hemifusion and has been proposed as a key step in LNP-mediated delivery, one of the possible endosomal escape mechanisms ^34,65–67^. Its rapid formation suggests that this formulation is intrinsically competent for engaging the endosomal membrane. However, due to the accelerated hydrophobic mixing inherent to Martini and the reduced accuracy of the CG model in forming pores^68,69^, the subsequent fusion pore-opening step could not be resolved reliably at the CG level, forcing us to change again from resolution model back-mapping the stalked configuration to atomistic resolution to follow the next stage of the pathway (see Methods).

**Fig. 3:**
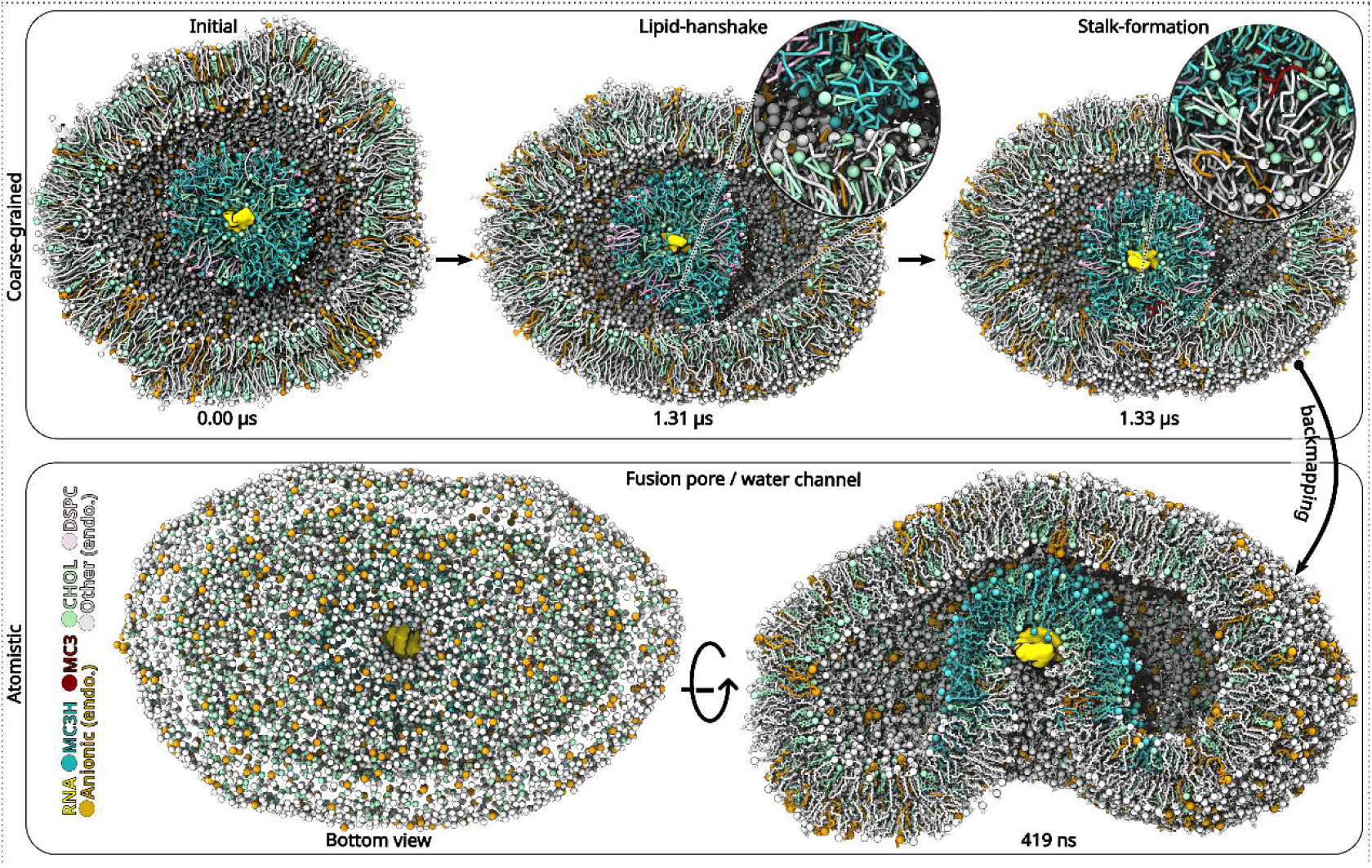
Molecular sequence of LNP–endosome engagement, hemifusion and pore formation. Top row (coarse-grained): successive representative snapshots show a naked (without PEG-lipid) HER2–siRNA LNP (taken from pH 6) contacting a prebuilt bilayer endosomal vesicle and progressing through early fusion intermediates. Initially (at t = 0), the particle is placed in the center of the bilayer vesicle. Within ∼ 1.3 µs, the LNP establish extensive lipid–lipid contacts (“lipid–handshake”) with the inner layer of the vesicle, and a shallow fusion stalk forms by t ∼ 1.33 µs (right). Insets: zoom into the contact region and highlight interdigitating lipid tails and the first lipidic continuity between LNP and endosomal leaflets. Bottom row (atomistic, backmapped): the CG stalk configuration was converted to an all-atom model and simulated further. The representative atomistic structure (bottom right) shows that within ∼ 419 ns of atomistic sampling, a water-filled fusion pore/channel nucleates at the stalk site and connects the hydrated LNP core to the endosomal lumen. The pore provides a physically plausible route for cargo access to the membrane opening. Color code used: RNA (gold), protonated MC3 (cyan), neutral MC3 (red), cholesterol (mint), DSPC (pink), PEG-lipid residues (gray), and anionic endosomal lipids (orange), other zwitterionic lipids of endosome (silver).

Remarkably, atomistic MD simulations started from the CG stalked conformation show the very fast (∼400–500 ns) formation of a distinct water-filled channel at the fusion interface (Fig. 3, bottom), connecting the hydrated LNP core with the endosomal lumen (<u>Movie SM6</u>). The fusion pore allows the siRNA to become partially exposed to the channel (Fig. 3, bottom view). This structure represents the second key intermediate of the endosomal escape process: a nascent fusion pore for nucleic acid release. It is important to emphasize that the fusion stalk and transient water-filled channel observed in our simulations should be interpreted as molecular events occurring within the specific MC3-based siRNA LNP model investigated here. However, these structures correspond to well-established intermediates in membrane fusion theory, where stalk formation and subsequent pore nucleation are considered key steps in membrane remodeling and fusion processes^70,71^. So, the process observed here may be valid for some other similar LNP formulations. Additionally, during the course of the MD simulation, no spontaneous release of the siRNA to cytoplasm is observed, which may correlate with the very low efficiency of endosome escape detected experimentally, as only ∼1–2% of internalized siRNA reaches the cytosol^15^, with some delivery processes taking place in a time scale of days (See experimental cytosolic delivery in <u>Fig. S1</u>) to months^46,72^.

### Energetic and structural determinants of siRNA escape from the endosomal pore

Since spontaneous RNA escape did not occur (<u>Movie SM6</u>) within the accessible atomistic timescales, we combined enhanced-sampling simulations with targeted structural analyses. To this end, we first generated a rough escape pathway by applying steered MD (SMD) forces to pull the RNA away (<u>Movie SM7</u> and see Methods) from the LNP core using the center-of-mass (COM) separation distance (r) between RNA and LNP as the collective variable (CV). This yielded an atomistic configuration of a translocated RNA threaded through the water-filled pore (Fig. 4a). The associated potential of mean force (PMF) was obtained by umbrella sampling (US) considered a fixed MC3 protonation states (that corresponds to pH=6 from above CpHMD ensemble). These calculations led to unrealistic free energy barriers (<u>Fig. S1</u>7), due to expected sampling limitations involving the strong stabilizing interactions of the protonated lipids (MC3H). These protonation states were assigned assuming an intact LNP structure (Fig. 2a), which may represent an artefact once structural rearrangements occur and the pore opening to the cytoplasm, where the pH is higher than in the endosome. Additionally, the presence of the water channel (Fig. 3, bottom) generates a different environment that can completely change the pKa of the ionizable lipids. This suggests that MC3 ionic state should be considered part of the reaction coordinate.

**Fig. 4:**
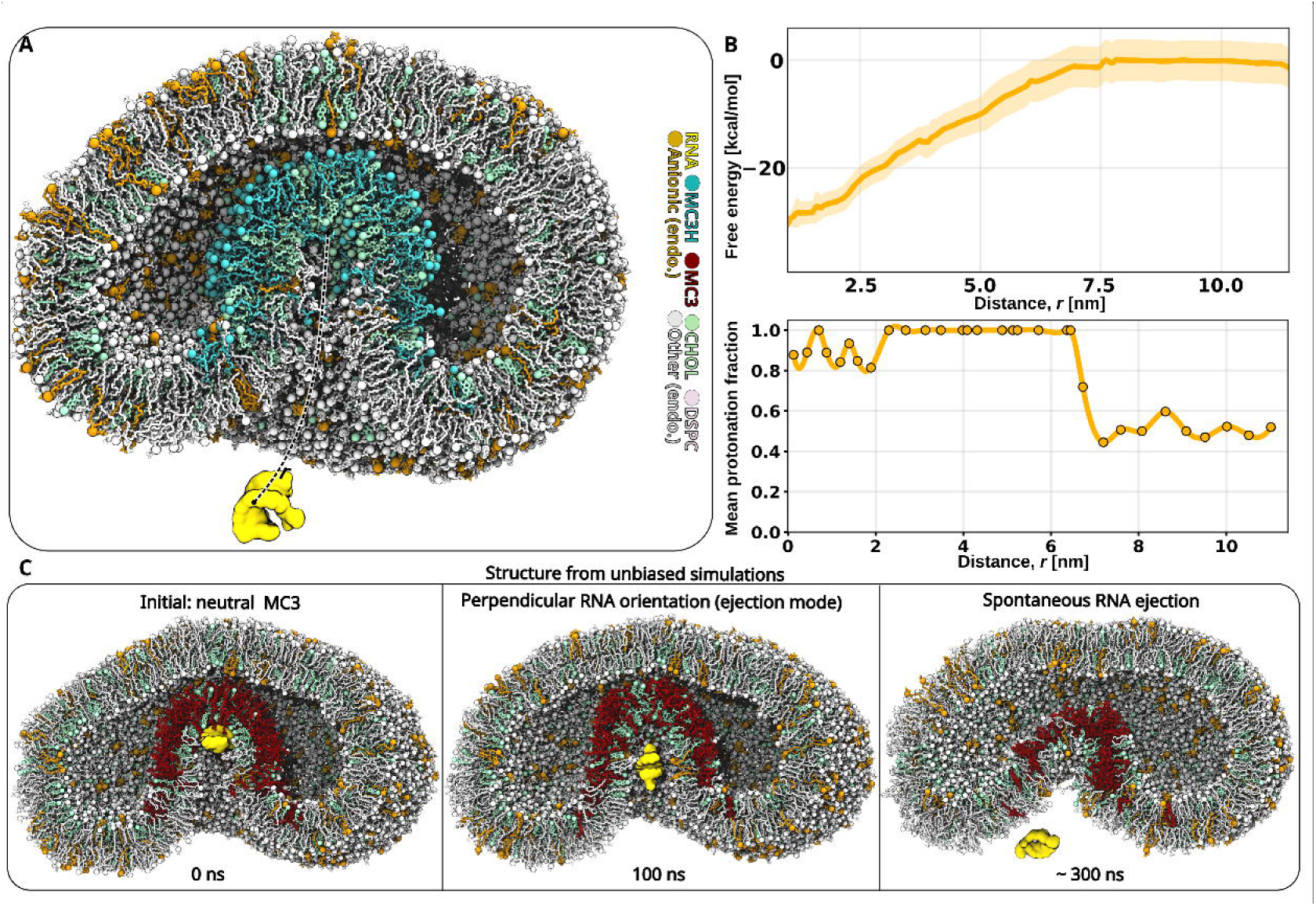
Energetics and atomistic pathway of siRNA translocation through the endosomal pore. (A) Representative atomistic snapshot of the RNA threaded through the fusion pore used as the initial configuration for biased sampling (center-of-mass pulling). Color code: RNA (gold), protonated MC3 (cyan), neutral MC3 (red), cholesterol (mint), DSPC (pink), PEG-lipids (gray), anionic endosomal lipids (orange), and other zwitterionic lipids of endosome (silver). The dashed line indicates the pulling coordinate (LNP COM → RNA COM distance, r) used as the collective variable for steered MD and umbrella-sampling. (B, top): Free-energy profile for RNA translocation computed with CpHMD– umbrella sampling (CpHMD–US) at endosomal pH (6.5). The PMF reports a substantial energetic barrier for pulling the RNA from the particle core, consistent with the low experimental endosomal escape efficiency^15^. Convergence of the free energy profile presented in <u>Fig. S1</u>8. Data are presented as mean ± SD. (B, bottom): Coupling between protonation and RNA position. Distance-resolved mean protonation fraction of MC3 residues computed on the umbrella-sampling windows. This map shows how local MC3 protonation changes as the RNA moves from the protected core into the pore and toward bulk solvent (mimicking cytosol). (C) Unbiased atomistic trajectories testing mechanistic hypotheses that the MC3 protonation-dependent electrostatic is the main deterministic factor behind LNP’s endosomal bottleneck. Representative unbiased simulations starting from the fusion-pore intermediate are shown. Left: Representative starting snapshot from the fusion-pore intermediate, but MC3 residues were converted to neutral (deprotonated) form. Middle: shows that within 100 ns, RNA adopted a perpendicular “ejection” orientation. Right: shows that RNA then spontaneously translocates through the pore within 300 ns.

To couple the MC3 ionic state and RNA escape, we combined US and CpHMD, and simulated at pH 6.5, allowing MC3 residues to change protonation dynamically. The resulting PMF (Fig. 4b, top) shows a reduced, but still substantial (∼29 kcal/mol) barrier for RNA translocation through the pore. This value, which seems reasonably well converged (<u>Fig. S18</u>) is consistent with the low efficiency of endosomal escape^15^, as well as our in-vitro experimental results that indicated endosomal escape in the days time-scale (see <u>Fig. S1</u>). Analysis of the trajectories show that MC3 lipids that contact RNA in the bound state remain highly protonated along the pulling pathway (Fig. 4b, bottom). Consequently, the siRNA remains tightly bound via electrostatic and water-mediated contacts to a cationic shell of MC3H even after the pore opens. As the apolar parts of the cationic lipids want to remain in an apolar environment, MC3H groups bridging RNA and the neutral lipid environment act as a break for the escape of the RNA. Thus, leaving the LNP implies a partial change in the cationic environment from MC3H to inorganic cations of the water solution (see Movie SM8).

Next, we tested two mechanistic ideas to justify a faster escape: (i) the reorientation of RNA that align its helical axis with the water pore to generate an “escape geometry”, as suggested in a previous study,^34^ and (ii) reduction of MC3 charge (deprotonation) as a way to reduce electrostatic MC3-RNA interactions and the slight higher pH of cytoplasm. To test the first idea, we manually reorient the RNA into a perpendicular orientation, but in all such trajectories, the RNA relaxed back to a surface-aligned configuration because of strong MC3H–phosphate attraction, and no spontaneous translocation was detected (<u>Movie SM9</u>). We then tested the second hypothesis by neutralizing all MC3 lipids (Fig. 4c, left). Almost immediately, the RNA readily adopted the perpendicular escape geometry (Fig. 4c, middle) and then spontaneously translocated through the fusion pore within the unbiased simulations (Fig. 4e, right; see also <u>Movie SM10</u>), suggesting a translocation efficiency much larger than that experimentally detected. The fully neutralized MC3 system should be interpreted as a proof-of-concept strategy designed to isolate the role of MC3–siRNA electrostatic interactions in cargo release. Although complete neutralization of all MC3 lipids is unlikely to occur in intracellular conditions, these results demonstrate that weakening such interactions strongly promotes siRNA translocation through the fusion pore. This finding suggests that local reductions in MC3 protonation near the RNA, even if partial, may facilitate cargo disengagement and thereby contribute to the release process.

Together, these results suggest that although early endosomal pH (∼6) is sufficient to drive protonation and membrane engagement, as charged MC3H interacts with the anionic endosomal lipids—favoring stalks and pore formation. Once a fusion pore opens, the interior of the LNP becomes directly exposed to the endosomal lumen and the near-neutral cytosolic environment, such that the local environment inside the pore becomes progressively more hydrated and shifted toward higher pH, facilitating changes in lipid protonation. A well-designed ionizable lipid should therefore rapidly deprotonate in response to this subtle local pH shift, weakening its affinity for RNA so that the cargo can escape. In our formulation, the unusually high local pKa of MC3 residues near the RNA prevents this final neutralization step, providing a mechanistic explanation for why endosomal escape remains slow and highlighting a clear design direction: future ionizable lipids must balance endosomal activity with sufficiently low local pKa near nucleic acids to permit disengagement once a fusion pore has formed.

In conclusion, this multiscale study establishes a coherent computational framework that links lipid composition, cargo electrostatics, and pH-dependent protonation to the structural organization and intracellular behavior of siRNA-loaded LNPs. By integrating coarse-grained modeling with atomistic constant-pH simulations, we demonstrate that siRNA is not merely a passive payload, but a key structural determinant that stabilizes a hydrated core, whose local organization is conserved across length scales despite differences in global morphology. This organization persists across a broad pH range and gives rise to a heterogeneous protonation landscape, in which local apparent pKₐ values in the vicinity of the RNA differ substantially from those at the particle periphery. Such spatially resolved protonation provides a mechanistic explanation for a long-standing paradox of LNPs: robust RNA encapsulation and near-neutral surface charge at physiological pH, yet inefficient cytosolic release following endosomal uptake.

While the present study focuses on MC3-based LNPs, the proposed mechanism is expected to extend to other LNP formulations. Clinically relevant ionizable lipids, such as ALC-0315 and SM-102, exhibit different apparent pKₐ values in LNPs, with ALC-0315 displaying a lower apparent pKₐ than MC3 and SM-102 a higher one^73,74^. Based on the mechanism proposed here, a higher apparent pKₐ is expected to increase the fraction of protonated ionizable lipids at a given endosomal pH, potentially enhancing lipid–RNA electrostatic interactions and favoring cargo retention. Conversely, a lower apparent pKₐ may reduce lipid protonation and facilitate earlier cargo release. However, these trends are likely to be modulated by the local molecular environment, and the resulting distribution of local pKₐ values may be as important as the average apparent pKₐ itself. Consequently, the quantitative behavior of different LNP formulations will depend not only on the intrinsic properties of the ionizable lipid, but also on its interactions with neighbouring lipids and the encapsulated RNA, making the balance between cargo retention and release highly formulation dependent.

Extending these insights to a model endosomal environment, our simulations support a physically plausible sequence of events in which LNPs engage the endosomal membrane, form a hemifusion stalk, and transiently open a water-filled pore connecting the LNP core to the endosomal lumen. However, even after pore formation, RNA release remains kinetically hindered by persistent electrostatic coupling to protonated ionizable lipids in the immediate RNA–lipid environment. This finding indicates that, for the formulation studied here, the dominant bottleneck in endosomal escape is not solely pore nucleation, but the inability of RNA-associated ionizable lipids to sufficiently deprotonate within the highly hydrated fusion interface. While alternative escape pathways—such as vesicle budding-and-collapse or size-dependent rupture mechanisms^75^ — have been proposed and may operate for other formulations or cargos, the present results demonstrate that local protonation chemistry alone can impose a strong kinetic barrier to release.

Taken together, these results emphasize a mechanistic principle rather than a formulation-specific outcome: effective endosomal escape requires not only protonation-driven membrane engagement, but also timely local deprotonation of ionizable lipids in the immediate vicinity of the nucleic acid once a permeation pathway has formed. Our simulations suggest that this process is governed by local, environment-dependent protonation behavior—rather than global apparent pKₐ values alone. In particular, the elevated local pKₐ values observed near the siRNA indicate that lipid structure can influence cargo retention, and may therefore be essential for understanding and ultimately overcoming the endosomal escape bottleneck in RNA delivery. From a formulation-design perspective, these findings suggest that optimizing the apparent pKₐ of an ionizable lipid may not be sufficient to ensure efficient cargo release. Instead, future ionizable lipids may benefit from maintaining sufficient protonation at low pH to promote endosomal membrane engagement while avoiding excessive stabilization of the protonated state in the immediate vicinity of the RNA at neutral, physiologic pH. This could potentially be achieved through modifications to its headgroup linker or other lipid features that modulate local protonation.

Further refinements of the model should account for size-dependent effects, dependence on cargo, role of corona proteins and other cellular or systemic effects ignored in our current simulations. Future multiscale simulations of mRNA-containing LNPs will also help determine whether the mechanistic principles identified here extend to the intracellular delivery of much larger RNA cargos or whether additional mechanisms emerge during mRNA transfection.

## Methods

### Coarse-grained (CG) model construction

#### Prebuilt large LNP model construction

The construction of the large prebuilt LNP was based on a protocol developed within the Martini 3 framework^76^ to construct inverse hexagonal LNPs^37^. Self-assembly was not used for this system due to its size and compositional complexity, making spontaneous assembly computationally prohibitive. In addition, experimental LNPs are kinetically assembled under rapid-mixing conditions, such that their morphology may reflect non-equilibrium assembly pathways rather than an equilibrium structure^59^. A conventional self-assembly simulation starting from randomly distributed components would therefore not reproduce the complete experimental formulation process. The pre-assembled protocol was therefore used to generate an initial structure consistent with the experimentally characterized internal organization of siRNA-loaded MC3 LNPs^77^.

The protocol for the construction of the full LNP aggregates several tools such as TS2CG^78^, Packmol^79^, VMD^80^ and the python packages, MDAnalysis^81^, and MDVContainment^82^ to facilitate the assembly of complex LNP structures. CG ionizable lipid, sterol and helper lipid parameters were obtained from the following Martini 3 publications^37,69,83^. The RNA model used is a 21 nucleotide-long polyA dsRNA sequence modeled using the same preliminary model employed in recent LNP and biomolecular condensate studies^84^. In this model, bead mapping and bonded parameters were based on the phosphate and ribose of the Martini 2 nucleic acid models^85,86^, with adaptations to the bead types. Phosphate groups are represented by Q5 beads, while ribose is represented by SN3a-SP1 fragments. Aromatic nucleobase rings were already available with Martini 3^76^. Since this model was still a prototype, the *π*-stacking between nucleobases was reinforced via an elastic networks.

The construction of the full LNP consists of a stepwise protocol, starting with the construction of a single cylinder coated with MC3H and cholesterol (ratio 56:43), using TS2CG^78^. The cylinder is then filled with the RNA cargo, water and ions using Packmol. This cylinder is then replicated in the z-axis to achieve a total length of 50 nm.

The following step consists in the replication of the channel in the x and y axis, and consequent identification of the different channels using MDVContainment^82^, and consequent extraction of a whole hexagon with 60 channels in total.

Once the LNP core was assembled, the coating was generated by first creating a triangulated surface of the hexagon using VMD^80^ in combination with Blender. This triangulated mesh was then used as input for TS2CG to build both a monolayer and a bilayer with a lipid composition of MC3H:CHOL:DSPC:DPG-PEG at 30.5:43.4:20:6 ratios. Since the channels at the top of the hexagon are open, a bilayer was placed at each channel end, with the hydrophilic headgroups oriented toward both the channel interior and the surrounding solvent. Along the length of the channels, a monolayer was applied instead, with the headgroups facing the solvent. This procedure yielded a coated LNP with an overall diameter of approximately 50 nm and a total lipid ratio of DSPC:CHOL:MC3H:DPG-PEG = 8.5:35.1:46.5:1.8.

After construction, the resulting LNP particle was solvated and Na⁺ and Cl⁻ ions were added to reach a physiological 150 mM salt concentration. Using GROMACS 2018.8^87^, and the Martini 3^76^ forcefield, the system was energy minimized in 500 steps, followed by 5 steps of equilibration, before 10 μs CG simulations in three replicates.

The target temperature was 310 K during both equilibration and production runs using, respectively, the Berendsen^88^ and Velocity-rescale thermostat^89^ with tau = 1 ps, coupling lipids separately from the solvent.

The pressure was maintained with an isotropic pressure coupling during relaxation and production run using respectively the Berendsen^88^ and Parinello-Rahman barostats^89,90^. Pressure was coupled with tau = 12 ps and the compressibility at 3×10^-4^ bar^-1^.

#### Self-assembly protocol for small LNPs

To examine whether the organization of MC3-based LNPs emerges naturally from the underlying RNA-lipid interactions, we generated a smaller, computationally tractable LNP by self-assembly rather than pre-building the structure. The goal was to let the particle form spontaneously and test whether the same internal local internal organization around RNA as in our large prebuilt model appears without imposing any initial geometry, consistent with previous CG self-assembly studies of nucleic-acid–loaded LNPs and related amphiphile systems^35,91^.

We used the same molar ratio as in our experimental formulation (DSPC:CHOL:MC3:DPG-PEG = 10:38.5:50:1.5), which is similar to the composition used in clinically approved MC3-based LNPs (e.g., Onpattro^13^). For the nucleic acid cargo, we included a short 10-bp HER2-targeting RNA duplex (sequence: 5’-CUCACAGAGA-3’), generated by trimming the unpaired terminal overhangs to avoid artifacts that can arise from excessive flexibility in CG RNA models.

A cubic box of 15.69 × 15.66 × 15.40 nm³ was used. All lipid and RNA molecules were inserted at random positions and orientations using the *gmx insert-molecules* module in GROMACS 2018.8^87^, with Martini3.0^76^ forcefield. Since MC3 has an intrinsic pKa of 9.47^51^ and LNP manufacturing occurs at acidic pH (∼4), we initialized all MC3 molecules in their protonated, cationic form (MC3H), as also done in previous computational studies of ionizable lipid assembly^25,92^.

During LNP production, lipids are typically dissolved in ethanol before rapid mixing with an aqueous buffer. To mimic this environment, we solvated our system in a 60:40 (water:ethanol) mixed solvent. Counterions (Na⁺ and Cl⁻) were added to neutralize the total charge and reach a physiological 150 mM salt concentration.

Three independent self-assembly replicas were prepared using different random seeds. Each system was minimized and equilibrated before running multi-microsecond CG simulations.

#### Prebuilt small LNP model construction

Similarly to the large LNP model, the construction of the small prebuilt LNP was based on a protocol developed within the Martini 3 framework^76^ to construct inverse hexagonal LNPs^37^. The construction of the full LNP consists of a stepwise protocol, as described above in section **1.1** of the materials and methods. The procedure yielded a coated LNP with an overall diameter of approximately 10 nm and a total lipid ratio of DSPC:CHOL:MC3H:DPG-PEG = 13.2:35.3:49.5:1.9.

After construction, the resulting LNP particle was solvated and Na⁺ and Cl⁻ ions were added to reach a physiological 150 mM salt concentration. Using GROMACS 2018.8^87^, and the Martini 3^76^ forcefield, the system was energy minimized in 500 steps, followed by 5 steps of equilibration, before 5 μs CG simulations in three replicates.

The target temperature was 310 K during both equilibration and production runs using, respectively, the Berendsen^88^ and Velocity-rescale thermostat^89^ with tau = 1 ps, coupling lipids separately from the solvent.

The pressure was maintained with an isotropic pressure coupling during relaxation and production run using respectively the Berendsen^88^ and Parinello-Rahman barostats^89,90^. Pressure was coupled with tau = 12 ps and the compressibility at 3×10^-4^ bar^-1^.

#### Endosomal model construction

To investigate RNA transfection from the small LNP, we constructed models of an endosomal membrane and a target plasma membrane using the TS2CG^78^ and INSANE^93^ tools.

In the endosomal transfection system, the endosomal membrane was generated with TS2CG^78^, which converts triangulated surfaces (TS) into coarse-grained membrane models suitable for molecular simulations. For this system, we used a spherical triangulated surface of 30 nm diameter and applied a lipid composition representative of the endosomal environment. (PLPC : SAPC : SDPC : PLPE : PDPE : SAPE : SAPI : SDPI : DSM : OSM : CHOL = 10:17:6:2:4:7:6:2:8:8:32). All CG lipid parameters were obtained from the new Martini3.0 lipidome paper^69^.

Following construction, the small LNP was placed at the center of the endosome, and counterions (Na⁺ and Cl⁻) were added to neutralize the total charge and reach a physiological 150 mM salt concentration. Using GROMACS 2018.8^87^, the system was energy minimized in 500 steps, followed by 5 steps of equilibration, before being simulated in six independent replicates, each with a duration of approximately 1.5 μs.

### Mapping and backmapping

To connect the atomistic description of the LNP with the longer-timescale coarse-grained (CG) simulations—and to ensure that the structural trends we observe are not artifacts of a single resolution—we routinely switched between atomistic and CG representations. The main goal was to get the best of both worlds. For converting atomistic structures to the Martini3.0 CG, we used the fast-forward mapping package^95^, which provides an automated and reproducible workflow for atomistic to CG transformations, using a customized atom to bead mapping file. This workflow was essential for reconstructing the CpHMD representative structures and running for a longer timescale in CG.

Backmapping from CG to atomistic was performed using CG2AT^96^. CG2AT uses a fragment-based reconstruction approach where chemically accurate atomistic fragments are placed onto CG bead positions using a library of customized “template fragments” that encode the correct stereochemistry, torsions, and connectivity. This method has been shown to outperform earlier interpolation-based backmapping schemes by producing physically plausible atomistic geometries that require only short equilibration^96–98^. In our workflow, CG2AT was essential for reconstructing the CG self-assembled LNPs prior to atomistic CpHMD. After backmapping, each system was energy-minimized and equilibrated for at least 100 ns to remove any structural artifacts inherited from the reconstruction.

### Molecular dynamics simulations

All molecular dynamics (MD) simulations were conducted in the isothermal–isobaric (NPT) ensemble using GROMACS 2018.8^87^. Coarse-grained (CG) and atomistic models were described with the Martini 3.0^76^ and CHARMM36^99^ force fields, respectively. The system temperature was maintained at 300 K through the v-rescale thermostat^89^, while the pressure was controlled at 1 bar using the isotropic C-rescale barostat^100^. Periodic boundary conditions were applied in all three spatial dimensions.

For electrostatics, reaction-field interactions^101^ were employed in the CG simulations, whereas long-range electrostatics in the atomistic systems were treated with the particle-mesh Ewald (PME)^102^ method. Real-space cutoffs of 1.1 nm (CG) and 1.2 nm (atomistic) were used. Van der Waals interactions were modeled with a Lennard–Jones potential, using the same cutoff values as above and a switching function starting at 1.0 nm.

In atomistic simulations, bond lengths within lipid and DNA molecules were constrained using P-LINCS^103^, and SETTLE^104^ was applied to maintain the geometry of water molecules. The leap-frog integrator was used for propagating the equations of motion, with a time step of 2 fs in atomistic simulations and 20 fs in the CG systems. Before starting production runs, each system underwent energy minimization followed by a five-stage equilibration protocol in which the integration time step was gradually increased to its final value.

### SAXS Calculation from prebuilt large LNP CG model

The SAXS profile of the large LNP was calculated on the last frame of the 10μs replicates, and computed using the martini_saxs method described elsewhere^105,106^, which follows the method available at https://github.com/fabianschuhmann/martini_saxs. In the present study, the scattering range up to q_max_=0.6 nm^-1^ was used. The 1D profile was obtained by radial averaging onto N_q_=25 uniformly spaced target q-values over the selected q-range, where N_q_ denotes the number of points in the output q-grid.

### Constant-pH molecular dynamics (CpHMD)

To capture the pH-dependent behavior of studied LNP systems—including both the siRNA-loaded and empty particles—we performed explicit-solvent constant-pH MD (CpHMD) simulations using the GROMACS-CpHMD beta implementation (version 2021)^107^. Our goal was to allow all MC3 lipids in the system to titrate dynamically so that protonation changes could respond naturally to the local environment, something that standard fixed-charge MD cannot capture.

The CpHMD method is based on λ-dynamics, where the protonation state of each titratable site is represented by a continuous λ coordinate. The end states λ = 0 and λ = 1 correspond to the fully protonated and deprotonated forms, respectively, and intermediate λ values represent alchemical mixing states used only for sampling (similar in spirit to free-energy perturbation techniques). The method allows the system to visit different protonation states with minimal added computational cost; the main overhead comes from additional PME electrostatics evaluations, which slow simulations by ∼20% compared to standard MD but do not scale with the number of titratable sites. A small biasing term is added internally to improve sampling of physical protonation states, but this does not affect the underlying free-energy landscape. Overall, this framework provides a practical and efficient way to include protonation equilibria directly in MD simulations.

Because MC3 is not part of the standard CpHMD residue library, we parametrized it following the workflow described in the pH-builder manual (https://gitlab.com/gromacs-constantph/phbuilder)^108^. Briefly, we simulated a single MC3 molecule in water, solvated with ions and a buffer particle, at its intrinsic pKa (9.47)^51^. From this reference system, we obtained an initial estimate of the dV/dλ correction potential using the helper scripts provided in pH-builder. We then performed ten independent 100-ns simulations to refine this parameter by further using inverse-Boltzmann reweighting. Successful calibration is indicated when the λ distribution as a histogram is flat at pH = pKa (see <u>Fig. S19</u>) and the energetic contributions from the empirical bias terms V_bias_ and V_pH_ vanish, leaving only the force-field (V_ff_) and correction terms (V_corr_), which should cancel out (see Ref:^108^ for details).

All CpHMD production simulations for the small LNPs were set up using pH-builder and run over a broad pH range (4–14) with all 300 MC3 lipids treated as titratable. For each pH, two independent replicas (∼300 ns each) were performed to ensure reliable convergence of protonation fractions before downstream CG backmapping and analysis.

### Enhanced-sampling for RNA translocation

To understand the final and functionally most critical step of delivery—how the siRNA actually disengages from the LNP once a fusion pore or water channel has formed—we combined steered MD (SMD), umbrella sampling (US), and constant-pH umbrella sampling (CpHMD-US).

We first performed SMD to generate a physically realistic “enforced-translocation” pathway in which the siRNA was gradually pulled away from the LNP core (see Movie SM7). The collective variable (CV) was defined as the center-of-mass (COM) distance between the RNA phosphate backbone and the MC3 headgroup region of the LNP. A moving harmonic potential with a force constant of 239.23 kcal mol^-1^ nm^-2^ was applied, and the resulting trajectory was used to extract a series of evenly spaced configurations to serve as starting structures for umbrella sampling windows.

Using these SMD-generated frames, we first carried out classical umbrella sampling^109^ with fixed protonation states to test whether a conventional approach would be sufficient to describe RNA escape energetics. Each window was restrained using the same COM-based CV, with harmonic force constants ranging from 59.80 to 837.32 kcal mol^-1^ nm^-2^, optimized based on trial histograms. Each window was simulated for ∼200 ns, and the first 25% of each trajectory was discarded as equilibration. Free-energy profiles were reconstructed using the weighted histogram analysis method (WHAM)^110^, with uncertainties estimated via bootstrapping while accounting for time-series correlations. This fixed-protonation setup produced a very high and poorly converged barrier (Fig. S#), clearly indicating that the detachment of siRNA from the particle is tightly coupled to changes in the protonation state of MC3—and therefore cannot be captured by conventional US.

To properly account for this coupling, we next performed combined US with constant-pH MD (CpHMD-US) at pH 6.5, representative of the early endosomal environment after pore formation. In this framework, each umbrella window samples not only the positional fluctuations of the RNA but also the protonation dynamics of every MC3 lipid through the λ-dynamics scheme. The same CV, window spacing, and harmonic constants were used as in classical US. All MC3 molecules were treated as titratable, and their protonation states were allowed to fluctuate throughout each simulation according to the instantaneous local environment. Free-energy profiles were again obtained using WHAM, ensuring a consistent comparison between classical US and CpHMD-US.

### Analysis

All structural and protonation analyses were performed using MDAnalysis^81,111^ together with custom Python scripts developed for this work. Results were averaged over the three simulation replicas for each system. The molecular movies are made using VMD^80^ and molywood^112^.

The radius of gyration (Rg), center-of-mass distances, and time series were computed directly through MDAnalysis. These quantities were used to assess particle compaction and structural convergence across simulation conditions. Radial distribution functions (RDFs) around the siRNA were computed using custom Python scripts built on top of MDAnalysis. In contrast to standard atom–atom RDFs, we evaluated distances from the center-of-mass (COM) of each interior RNA molecule to the selected species (water, ions, or lipid beads) in every frame. For each configuration, distances were binned into spherical shells, averaged over all RNA molecules and post-equilibration frames. Water density was computed as a function of distance from the LNP COM by counting water beads in spherical shells. These profiles were used to quantify hydration changes across pH and between empty and RNA-loaded particles. Radial charge density ρ(*r*) and cumulative charge Q(*r*) were computed by binning atoms or beads into spherical shells centered on the LNP COM. Shell volumes were computed analytically, and charges were accumulated over the trajectory. Separate profiles were generated for: (i) lipids only, (ii) lipids + RNA, and (iii) full system including water and ions. These profiles were used to compare with experimental ζ-potential measurements.

For each titratable MC3 lipid, the protonation state was determined from the λ-coordinate. Frames with λ < 0.25 (since we always start the setup with protonated form) were assigned as protonated (MC3H) and frames with λ > 0.75 as deprotonated (MC3). Mean protonation fractions at a given pH were obtained by averaging over all MC3 residues and the full trajectory for each replica. For the distance-resolved analysis, MC3 molecules were binned according to their instantaneous distance from the RNA center of mass (or LNP COM for empty particles), and protonation fractions were computed separately in each bin. Titration curves (protonation fraction vs pH) were generated either globally (all MC3) or (locally) for each radial bin. The apparent pKa values were obtained by fitting to the Henderson–Hasselbalch equation. This approach was applied to both siRNA-loaded and empty LNPs.

## Experimental

### Lipids and nucleic acids

DLin-MC3-DMA [(6Z,9Z,28Z,31Z)-heptatriaconta-6,9,28,31-tetraen-19-yl-4-(dimethyl amino)-butanoate] was purchased from Tebubio; cholesterol from Merck; DPG-PEG (1,2-dipalmitoyl-rac-glycero-3-methylpolyoxyethylene-2000) from NOF America Corporation; and DSPC (1,2-distearoyl-sn-glycero-3-phosphocholine) from Avanti Lipids. The unmodified siRNA targeting HER2 (5’ AAG CCU CAC AGA GAU CUU GdTdT 3’) was obtained from Integrated DNA Technologies and the 5’-Cy5-labeled version of the same siRNA was sourced from Metabion.

### Lipid nanoparticle preparation

DSPC, DLin-MC3-DMA, cholesterol and DPG-PEG were dissolved in ethanol to prepare 10 mg/mL stock solutions. siRNAs were prepared by incubating equimolar amounts of guide and sense strands in annealing buffer (100 mM potassium acetate, 30 mM HEPES-KOH pH 7.4, 2 mM magnesium chloride) at 95 °C for 5 min, followed by 1 h at 37 °C. To formulate siRNA-loaded lipid nanoparticles, the annealing buffer in the siRNA solution was exchanged with 20 mM citrate buffer (pH 4) using an Amicon centrifugal filter (Merck Millipore, 3 kDa molecular weight cut-off). A lipid mixture (10:50:38.5:1.5 molar ratio of DSPC:DLin-MC3-DMA:cholesterol:DPG-PEG) was prepared in ethanol. The lipid mixture was added dropwise to the siRNA solution and resulting mixture (3:2 [vol/vol] citrate buffer:ethanol, 1:16.7 [wt/wt] siRNA:lipids) was incubated at 65 °C for 1 h under vigorous stirring (1600 rpm). The preparation was then sequentially extruded through 400 nm and 100 nm pore-size polycarbonate membranes (minimum 31 passes each) using an extruder (Avanti Lipids) coupled to a heating block at 65 °C. Finally, the LNPs were dialyzed against citrate buffer (20 mM citric acid, 20 mM sodium citrate, pH 4) for 2 h (7 kDa molecular weight cut-off) to remove ethanol, followed by HBS buffer (20 mM HEPES, 20 mM NaCl, pH 7.4) overnight (7 kDa molecular weight cut-off) to neutralize the ionizable lipid.

### Lipid nanoparticle characterization

The average size and polydispersity index of the LNPs were determined by dynamic light scattering using a Zetasizer Nano (Malvern Instruments) with Zetasizer 8.01 software. A 1:10 dilution of the LNPs in 20 mM HBS buffer (pH 7.4) was analyzed in a low-volume quartz cuvette, with the following settings: material refractive index 1.4, absorption 0.001, dispersant refractive index 1.33, dispersant viscosity 0.8872 cP, and a 173° measurement angle. All samples were analyzed in triplicate with 12-15 runs per replicate. The surface charge of the LNPs was determined by electrophoretic light scattering using a a Zetasizer Nano with Zetasizer 8.01 software. A 1:100 dilution of the LNPs in 20 mM HBS buffer (pH 7.4) was transferred to a folded capillary cell and the analysis was conducted with the following settings: material refractive index 1.4, absorption 0.001, dispersant refractive index 1.33, dispersant viscosity 0.8872 cP, and dielectric constant 75.5. All samples were analyzed in triplicate with up to 100 runs per replicate.

### Encapsulation efficiency

siRNA encapsulation efficiency was determined using a Qubit 4 fluorimeter and a microRNA assay kit (Thermo Fisher Scientific) for short RNA quantitation. siRNA concentrations were calculated from a standard curve generated using standard samples of known concentration, with the assay reagent binding to siRNA and producing a fluorescent signal proportional to its concentration. A 1:5 dilution of the LNPs in 20 mM HBS buffer (pH 7.4) was added to the assay reagent to determine unencapsulated siRNA concentration, and a 1:5 dilution of the LNPs in 0.1% Triton X 100 in 20 mM HBS buffer (pH 7.4) was used to determine total siRNA concentration. Encapsulated siRNA concentration was calculated by subtracting the unencapsulated siRNA concentration from the total siRNA concentration. Encapsulation efficiency was defined as the ratio of encapsulated siRNA to total siRNA and was expressed as a percentage.

### Cell culture

SKBR3 cells were obtained from ATCC and were cultured in McCoy’s 5A medium (Gibco) supplemented with 10% fetal bovine serum, 100 U/mL penicillin and 100 µg/mL streptomycin. Cells were maintained at 37 °C in a humidified atmosphere with 5% CO2.

### Cellular internalization assays

Cellular uptake of siRNA loaded LNPs was analyzed by fluorescence confocal microscopy. Cells were seeded on 24-well plates with 12 mm cover glasses on the well bottom at a density of 50000 cells per well in complete media and incubated at 37 °C in a humidified atmosphere with 5% CO2. Following overnight culture, cells were transfected with the LNP formulation at different doses (0.5 mL total volume) or left untreated as a control. After a 72 h incubation period, cells were fixed with 4% paraformaldehyde at pH 7.4 for 10 min at room temperature, permeabilized with 0.5% Triton X-100 in PBS for 5 min, and incubated in 1 µg/mL Hoechst 33342 for 5 min for nuclear staining. The cover slides were then isolated and mounted on microscope slides with Fluoromount-G (Electron Microscopy Sciences) and incubated overnight at 4 °C before imaging. The fluorescence of labeled nuclei and the Cy5 labeled siRNA was observed using an SPE confocal microscope (Leica) with LAS AF software. Hoechst 33342 fluorescence (461 nm emission) was visualized using a 405 nm laser diode (10% intensity, 1000 gain) and Cy5 fluorescence (667 nm emission) was visualized using a 635 nm laser diode (15% intensity, 1000 gain). The images were analyzed using ImageJ software (NIH software).

## Data Availability

All data supporting the findings of this study, including molecular dynamics trajectories, initial molecular configurations, force-field parameters, and analysis scripts, are publicly available in the Zenodo repository (DOI: <u>10.5281/zenodo.23419202</u> [https://doi.org/10.5281/zenodo.23419202]). The molecular dynamics trajectories are available through the MDDB platform, where they can be directly accessed and analyzed via https://mdposit.mddbr.eu/#/browse?option=lnp01. Source data underlying all figures are provided with this paper.

## Code Availability

Analysis scripts, are publicly available on Zenodo [https://doi.org/10.5281/zenodo.23419202].

## Acknowledgments

P.C.T.S and M.V. acknowledge the support of the French National Center for Scientific Research (CNRS). They acknowledge the support of the PSMN (Pôle Scientifique de Modélisation Numérique) and the Centre Blaise Pascal’s IT test platform at ENS de Lyon (Lyon, France) for the computer facilities. The platform operates the SIDUS solution (471) developed by Emmanuel Quemener.

## Author contribution

KAH contributed to the design of the study, performed all simulations, analyzed the results, and generated many of the figures and videos. MV assembled and tested the largest systems considered, co-designed parts of the analysis, and prepared several figures and videos. MV and PCTS also developed and generated all coarse-grained lipid and RNA models used in the study. PM performed and analyzed all experimental results. PCTS co-designed and supervised theoretical work and conceived the analysis of the results. MO conceived and co-designed the study and analysis and supervised both the experimental and theoretical work. All authors contributed to writing the manuscript.

