## Supplementary information for "pH-driven dynamics and mechanisms of LNP-mediated siRNA delivery"

| Reliability and reproducibility checklist for molecular dynamics simulations | Yes | N/A | Response |
|---|---|---|---|
| **1. Convergence of simulations and analysis** | | | |
| 1a. Is an evaluation presented in the text to show that the property being measured has equilibrated in the simulations (e.g., time-course analysis)? | ✓ | | Fig. 1, Fig. 2, Fig. S17, and Fig. S18 |
| 1b. Then, is it described in the text how simulations are split into equilibration and production runs and how much data were analyzed from production runs? | ✓ | | See the Method section |
| 1c. Are there at least 3 simulations per simulation condition with statistical analysis? | ✓ | | The equilibrium MD simulations were performed in triplicate. CpHMD and US were performed in two replicas considering their computational expense. |
| 1d. Is evidence provided in the text that the simulation results presented are independent of initial configuration? | ✓ | | SI figures and videos |
| **2. Connection to experiments** | | | |
| 2a. Are calculations provided that can connect to experiments (e.g., loss or gain in function from mutagenesis, binding assays, NMR chemical shifts, J-couplings, SAXS curves, interaction distances or FRET distances, structure factors, diffusion coefficients, bulk modulus and other mechanical properties, etc.)? | | ✓ | |
| **3. Method choice** | | | |
| 3a. Do simulations contain membranes, membrane proteins, intrinsically disordered proteins, glycans, nucleic acids, polymers, or cryptic ligand binding? | ✓ | | Nucleic acid, lipids and PEG |
| 3b. Is it described in the text whether the accuracy of the chosen model(s) is sufficient to address the question(s) under investigation (e.g., all-atom vs. coarse-grained models, fixed charge vs. polarizable force fields, implicit vs. explicit solvent or membrane, specific force field and water model, etc.? | ✓ | | All the validation and disclaimers are explicitly mentioned in the text. |
| 3c. Is the timescale of the event(s) under investigation beyond the brute-force MD simulation timescale in this study that enhanced sampling methods are needed? | ✓ | | The parameters and convergence criteria for the enhanced sampling method clearly stated in the text and showed in figures. |

| 4. Code and reproducibility | | | |
|---|---|---|---|
| 4a. Is a table provided describing the system setup that includes simulation box dimensions, total number of atoms, number of water molecules, salt concentration, lipid composition (number of molecules and type)? | | ✓ | Box dimensions and total number of atoms are mentioned in the respective subsection in Methods. |
| 4b. Are other parameters for the system setup described in the text, such as protonation state, type of structural restraints if applied, nonbonded cutoff, thermostat and barostat, etc.? | ✓ | | All the parameters are described in the text. |
| 4c. Is it described in the text what simulation and analysis software and which versions are used? | ✓ | | See the Methods |
| 4d. Are initial coordinate and simulation input files and a coordinate file of the final output provided as supplementary files or in a public repository? | | ✓ | Will be made available soon (upon acceptance for publication) |
| 4e. Is there custom code or custom force field parameters? | | ✓ | |

**Movie SM1:** Coarse-grained simulation of the large (~70 nm) siRNA-loaded LNP model demonstrating compaction into an inverted-hexagonal (HII) morphology.
**Movie SM2:** Spontaneous coarse-grained self-assembly of the small siRNA-loaded LNP (~10 nm) from randomly distributed lipids and siRNA.
**Movie SM3:** Atomistic equilibration of the backmapped small siRNA-loaded LNP model.
**Movie SM4:** Coarse-grained self-assembly and internal structural organization of the empty LNP (without siRNA).
**Movie SM5:** Coarse-grained simulation of naked LNP interactions with the endosomal membrane leading to stalk formation.
**Movie SM6:** Atomistic simulation showing the nucleation and opening of a water-filled fusion pore at the LNP–endosome interface.
**Movie SM7:** Steered molecular dynamics (SMD) simulation illustrating enforced siRNA translocation through the endosomal pore.
**Movie SM8:** Atomistic trajectory showing the persistent electrostatic coupling between siRNA and protonated MC3 lipids during pore engagement.
**Movie SM9:** Unbiased atomistic simulation testing RNA reorientation within the fusion pore under protonated MC3 conditions.
**Movie SM10:** Unbiased atomistic simulation demonstrating spontaneous siRNA ejection and translocation through the fusion pore following MC3 charge neutralization.

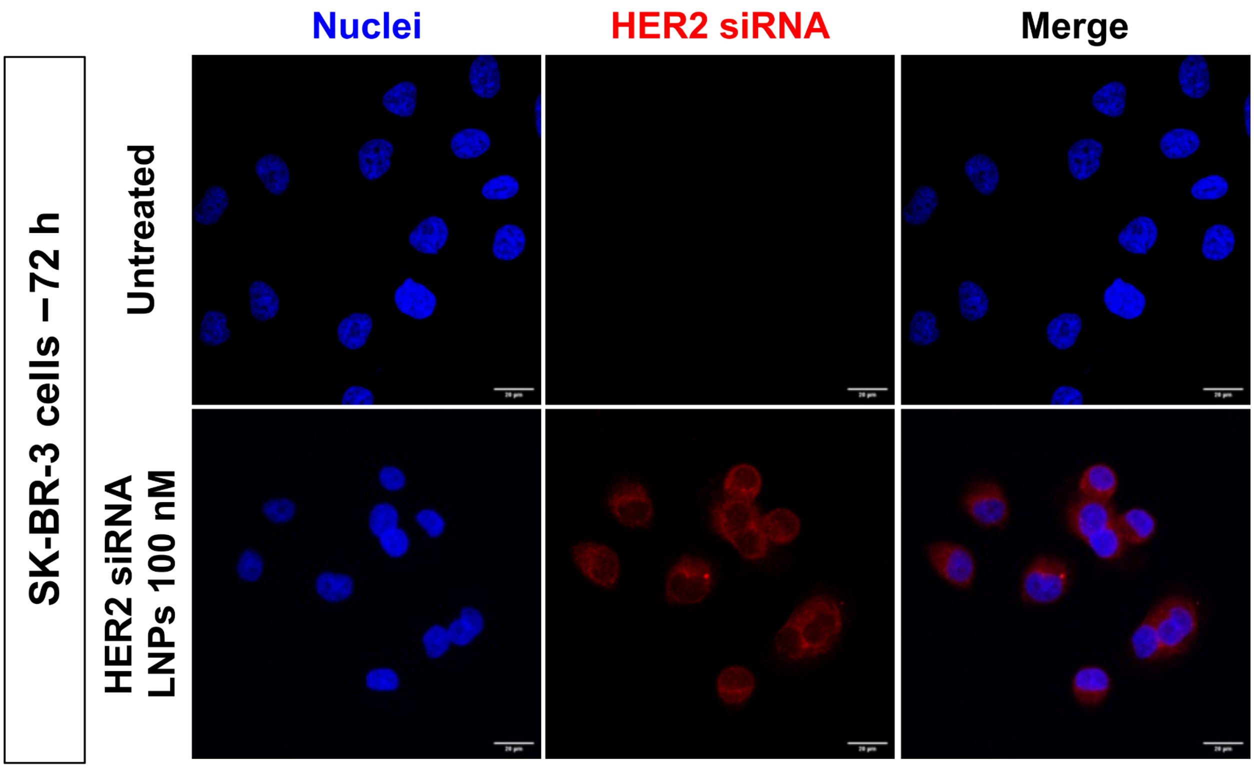


***Fig. S1*: Experimental validation of cytosolic delivery.**
*Fluorescence micrographs of SK-BR-3 cells were acquired after 72 h incubation with HER2-siRNA LNPs (100 nM). Nuclei (blue) and fluorescently labelled HER2 siRNA (red) are shown; the* ***top row*** *corresponds to untreated controls, and the* ***bottom row*** *shows LNP-treated cells. The intracellular red signal in treated cells confirms that a fraction of the siRNA reaches accessible intracellular compartments in vitro. Images are representative of three independent experiments.*

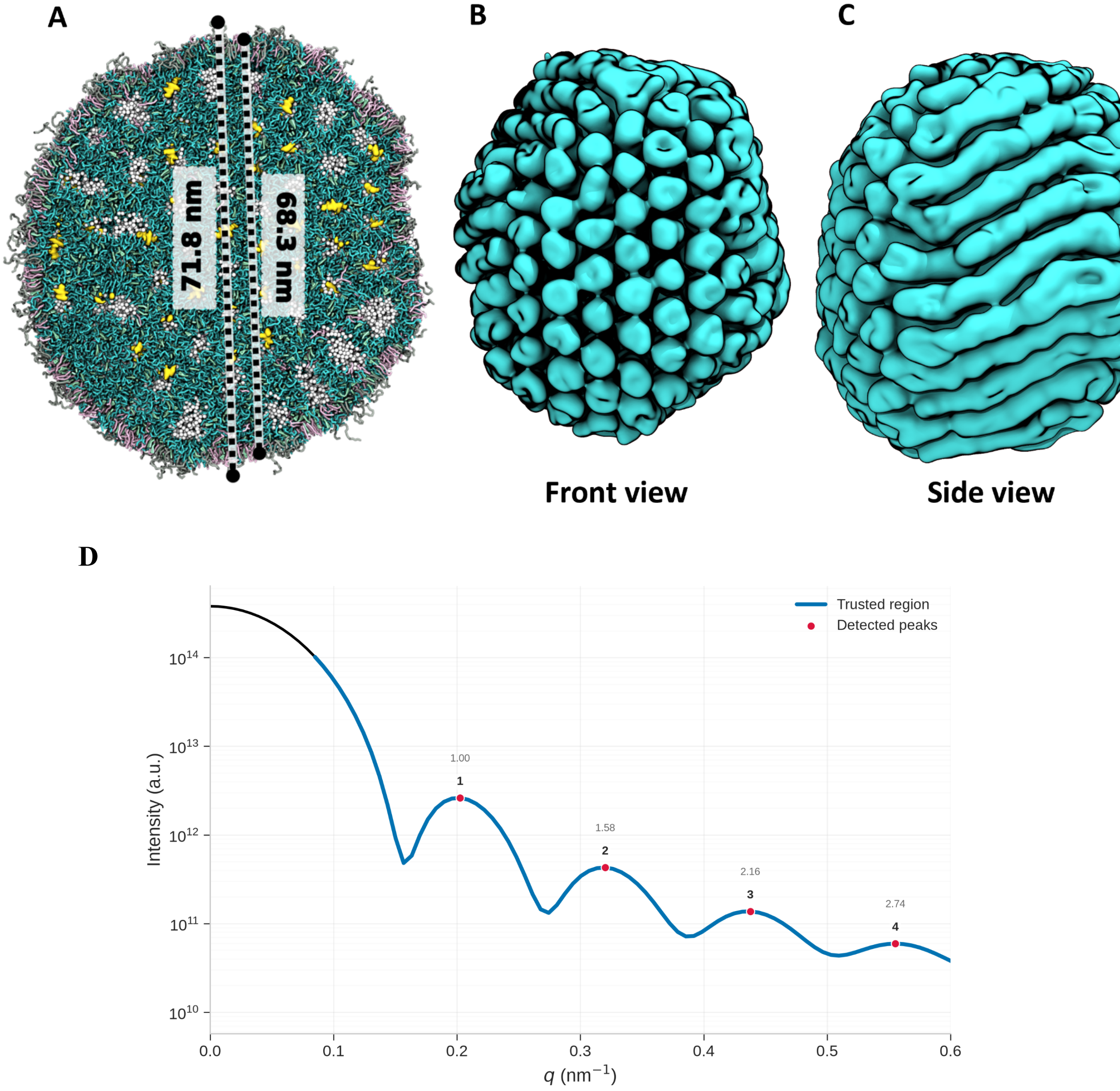


***Fig. S2*: Diameter and structural features of the large LNP model.**

*(A) The LNP diameter was calculated from the last 10 µs of the simulxation trajectories and defined as twice the maximum distance of any lipid bead from the center of mass (COM). Two cases are shown: including PEG-lipids (71.8 nm) and excluding PEG-lipids (68.3 nm). (B) Front view illustrating the inverted hexagonal ($H_{II}$) arrangement of RNA-containing channels within the LNP. (C) Side view of the channels formed by MC3 lipids. In panels (B) and (C), the structure is rendered using the QuickSurf representation in VMD based exclusively on the MC3 lipid NP bead group and shown in cyan. All images correspond to the final simulation frame at 10 µs. (D) SAXS profile calculated after 10 µs. The observed peak positions show approximate agreement with the characteristic reflections of a two-dimensional hexagonal lattice, providing evidence for hexagonal ordering.*

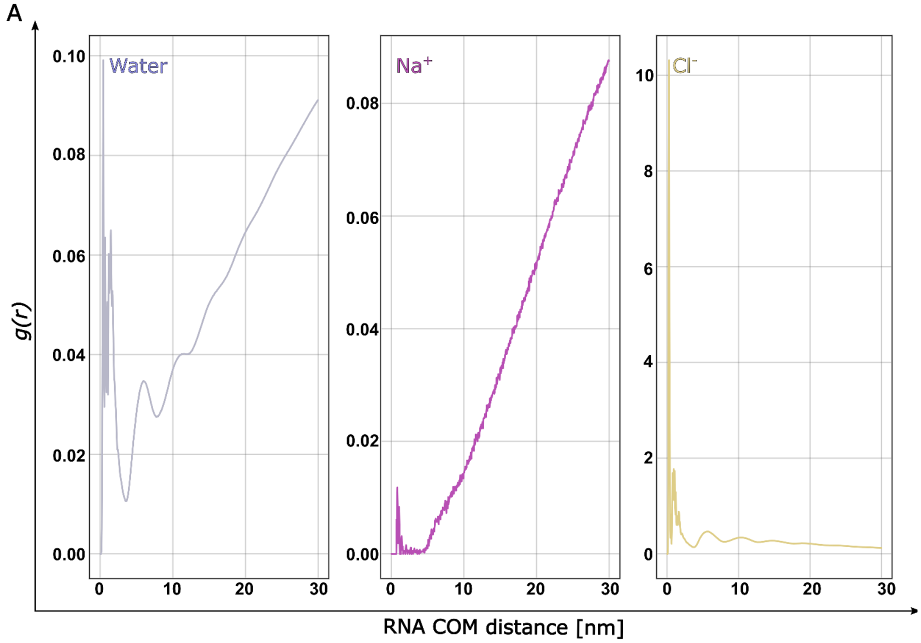


***[Fig. S3](#)*****: Radial distribution functions (RDFs) of water, $Na^{+}$, and $Cl^{-}$ beads (left to right), computed from the large (~70 nm) LNP model and averaged over two CG replicas.**

*(**A**) RDFs were calculated in the same way as the lipid–RNA RDFs in Fig. 1d (right panel), using the siRNA center of mass (COM) as the reference. The water profile shows that the LNP interior remains hydrated, forming several water channels that contain the inverted-hexagonal ($H_{II}$) organization seen in Fig. 1. $Na^{+}$ counterions show a small peak close to the RNA, reflecting partial screening of the phosphate backbone. However, because the LNP contains a large number of positively charged MC3H lipids, $Na^{+}$ is largely repelled from the particle interior and accumulates mostly outside the LNP surface. In contrast, $Cl^{-}$ counterions show a strong peak at short distances, as they preferentially interact with MC3H-rich regions, and their density gradually decreases at larger radii.*

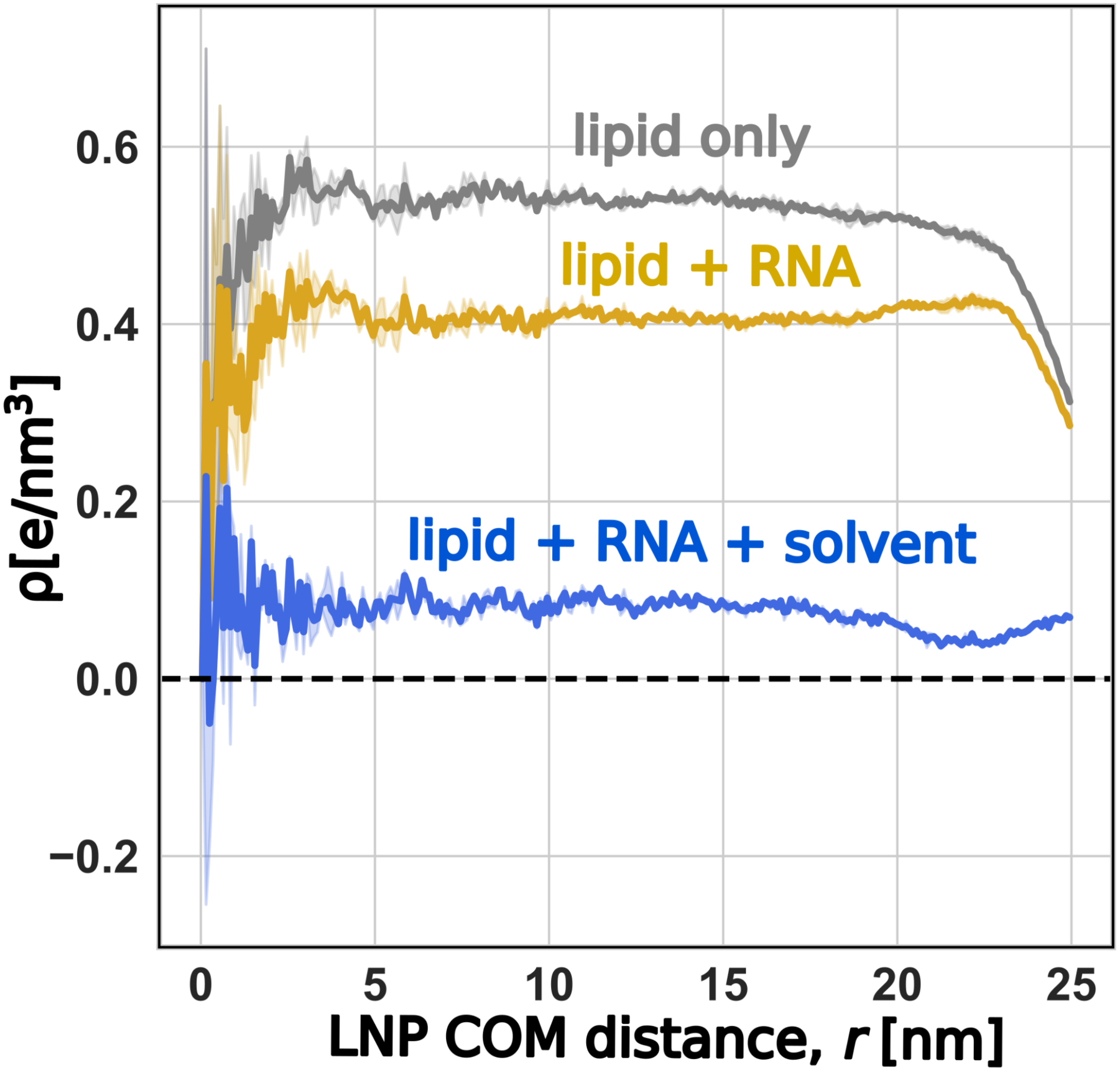


***Fig. S4*: Radial charge-density profiles of the large (~70 nm) LNP as a function of distance from the LNP center of mass (COM).**

*Charge densities were computed from coarse-grained trajectories (averaged over two replicas) by binning all charged beads into spherical shells around the LNP COM and normalizing by shell volume (see Methods). Three profiles are shown: lipids only, lipids + RNA, and lipids + RNA + solvent ions. The lipid-only curve exhibits a strongly positive charge density within the inner ~20 nm, reflecting the high abundance of protonated MC3H. Upon including RNA, the profile shifts toward neutrality due to the contribution of the siRNA phosphate backbone, partially compensating the MC3H charge. After adding $Na^+$ and $Cl^-$ (full system), the profile becomes nearly neutral throughout the particle, demonstrating that counterions efficiently screen the internal electrostatics.*

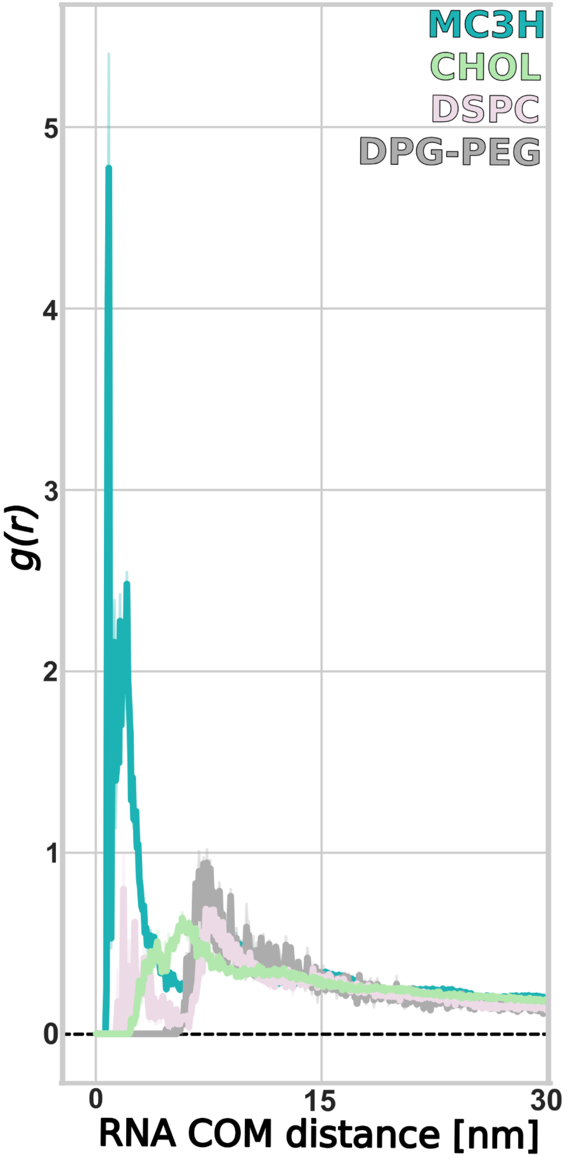


***Fig. S5*****: Radial distribution functions (RDFs) of lipid headgroups around a single siRNA molecule in the large (~70 nm) LNP model.**

*To allow a direct, like-for-like comparison with the self-assembled small LNP (Fig. 1e, right), we recomputed the lipid–RNA RDFs from the large particle by selecting only one representative siRNA and calculating lipid bead densities as a function of distance from its center of mass (COM). The profiles show the expected enrichment of protonated MC3H within the inner ~3 nm, reflecting strong electrostatic association with the RNA backbone, followed by broader distributions of CHOL, whereas DSPC and DPG-PEG are located near the LNP surface, consistent with their structural roles in packing and surface stabilization. Despite the greater heterogeneity and size of the large LNP, the qualitative trends closely match those of the small LNP, further validating that the reduced model preserves the same local lipid–RNA organization.*

.

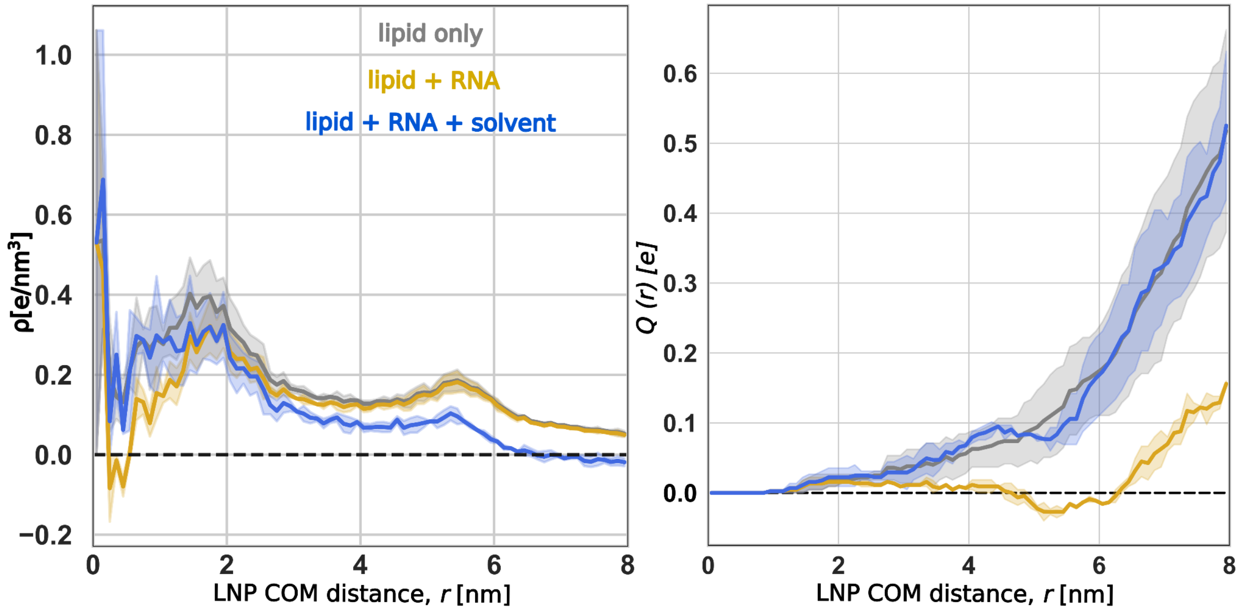


***Fig. S6*: Radial charge–density and cumulative charge profiles for the small self-assembled LNP.**
*(**Left**) Charge density ρ(r) (e·nm$^{-3}$) plotted as a function of distance from the LNP center of mass (COM) for three components: lipids only, lipids + RNA, and all species (lipids + RNA + solvent ions). The qualitative behavior closely matches that of the large LNP (Fig. S4): the "all-species" curve remains close to zero, demonstrating that counterions efficiently screen the combined lipid–RNA charge and the nanoparticle surface remains globally electroneutral. (**Right**) The corresponding cumulative charge Q(r), obtained by integrating ρ(r) from the center outward. In contrast to the large LNP, the lipid + RNA curve in the small self-assembled particle approaches near-zero and even slightly negative values at intermediate radii. In the small LNP, the siRNA might occupy a much larger fractional volume of the core, and its phosphates cluster closer to the geometric center relative to MC3H headgroups, which preferentially populate the inner–mid shell rather than the very core. When integrating charge from the center outward, the calculation therefore encounters a disproportionately high density of RNA anionic phosphate groups before sampling the compensating cationic MC3H layer, producing an apparent transient negative Q(r).*

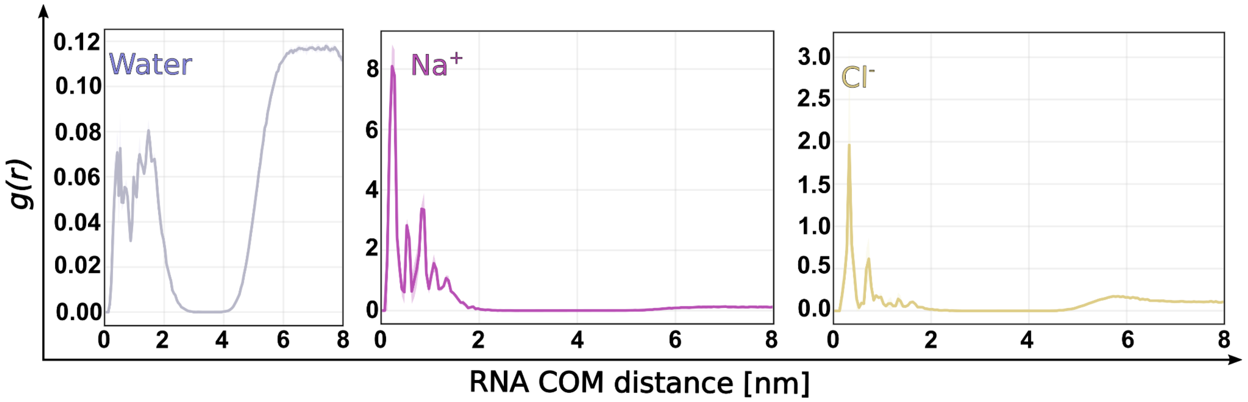


***Fig. S7: Radial distribution functions (RDFs) of water, $Na^+$, and $Cl^-$ beads (left to right) computed from the small self-assembled LNP, using the siRNA center of mass (COM) as the reference****.*

*The water profile shows that the LNP core remains hydrated and then gradually dehydrates toward the mid-radius before rising again near the particle surface as it approaches bulk solvent. $Na^+$ displays a sharp peak close to the RNA, reflecting its role in screening the phosphate backbone, but its density becomes very low beyond this region because of the presence of positively charged MC3H. In contrast, $Cl^-$ exhibits two preferred regions: one near the core, where it interacts with the MC3H lipids that cluster around the RNA, and another shallow peak toward the particle surface, where additional MC3H headgroups are exposed in the outer shell. Together, these profiles show that the small LNP reproduces the similar hydration and counterion organization trends as the larger model.*

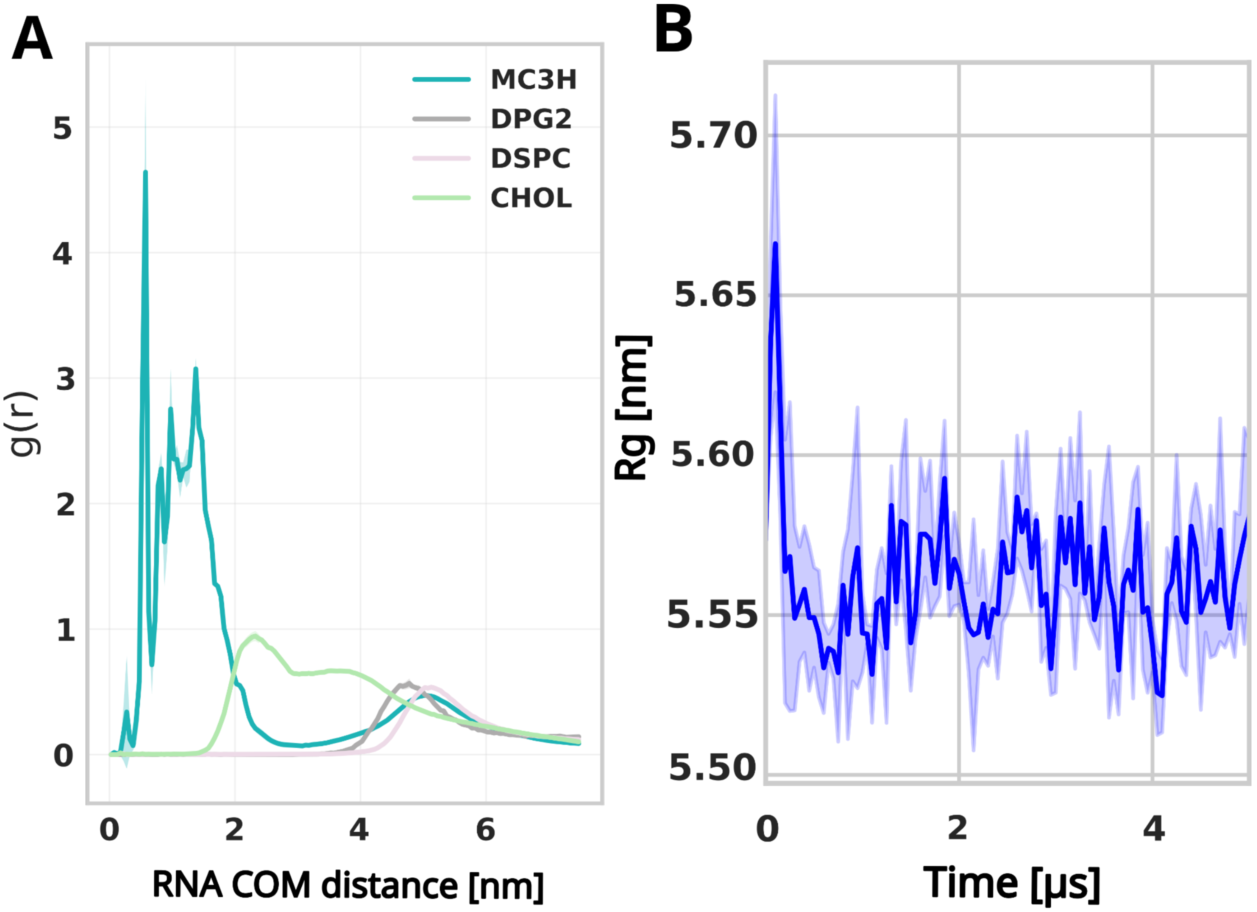


***Fig. S8: Structural organization of the small prebuilt LNP model.***
***(A)*** *g(r) of lipid headgroups around the RNA COM for the small prebuilt LNP particle. When compared with the self-assembled LNP (Fig. 1F), the same spatial ordering (MC3H core, cholesterol/intermediate, DSPC and PEG outer shell) is preserved.* ***(B)*** *Rg time series for the small prebuilt LNP particle (mean ± SD).*

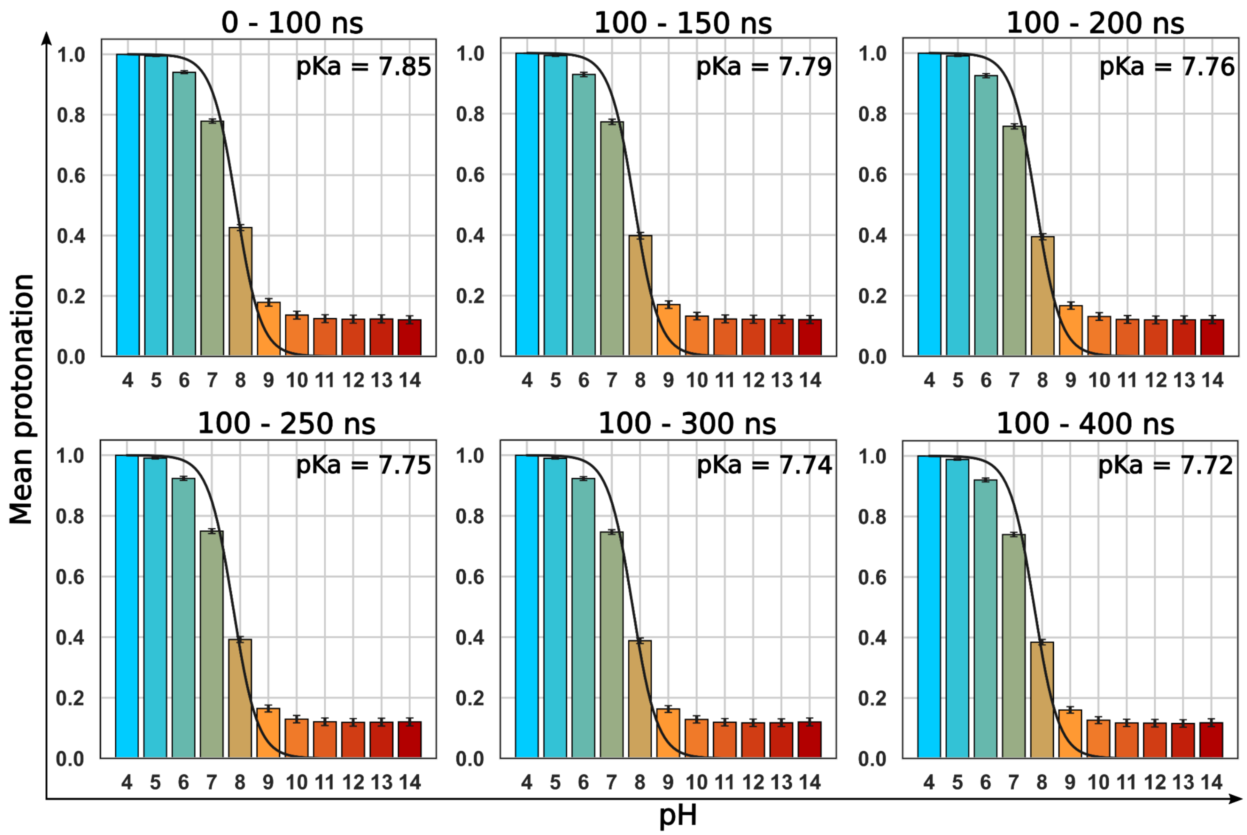


***Fig. S9: Convergence of MC3 protonation behavior in atomistic CpHMD simulations.***
*Mean protonation fraction of MC3 residues at each pH value (4–14) shown together with the corresponding titration curve fitted using the Henderson–Hasselbalch equation, calculated over two replicas and at time intervals of 0–100 ns, 100–150 ns, 100–200 ns, 100–250 ns, 100–300 ns, and 100–400 ns—used to assess convergence. Across these windows, the estimated pKa values remain stable within a narrow range, indicating that protonation equilibria of MC3 have essentially converged within the first ~300 ns of sampling. Extending the sampling to longer windows does not introduce systematic drift, supporting that the global titration curve reported in the main text (Fig. 2b) is statistically reliable.*

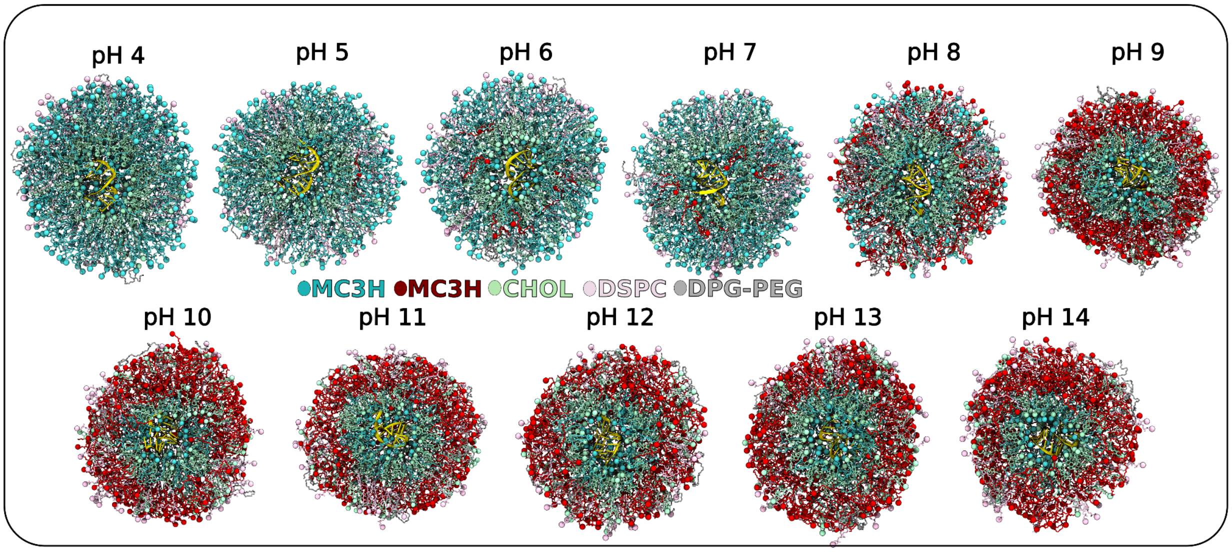


***Fig. S10: Representative atomistic snapshots of the siRNA-loaded LNP at each studied pH (4–14) from CpHMD simulations.***

*Shown are equilibrated configurations extracted from the CpHMD trajectories, illustrating the pH-dependent structural organization of the LNP. Color code: RNA (gold), protonated MC3H (cyan), neutral MC3 (red), CHOL (mint), DSPC (pink), DPG-PEG (gray). At acidic pH (4–6), the particle remains highly protonated (MC3H-rich) and adopts a more open, hydrated, and slightly elongated morphology. As pH increases (7–9), progressive MC3 deprotonation leads to compaction of the particle and formation of a more defined, spherical hydrophobic core. At alkaline pH (10–14), peripheral MC3 is largely deprotonated and the particle exhibits its most compact and ordered structure. These snapshots complement the quantitative analysis in Fig. 2 by visually highlighting the pronounced pH-driven transition from a loose, porous architecture at low pH to a tighter, more dehydrated organization at higher pH.*

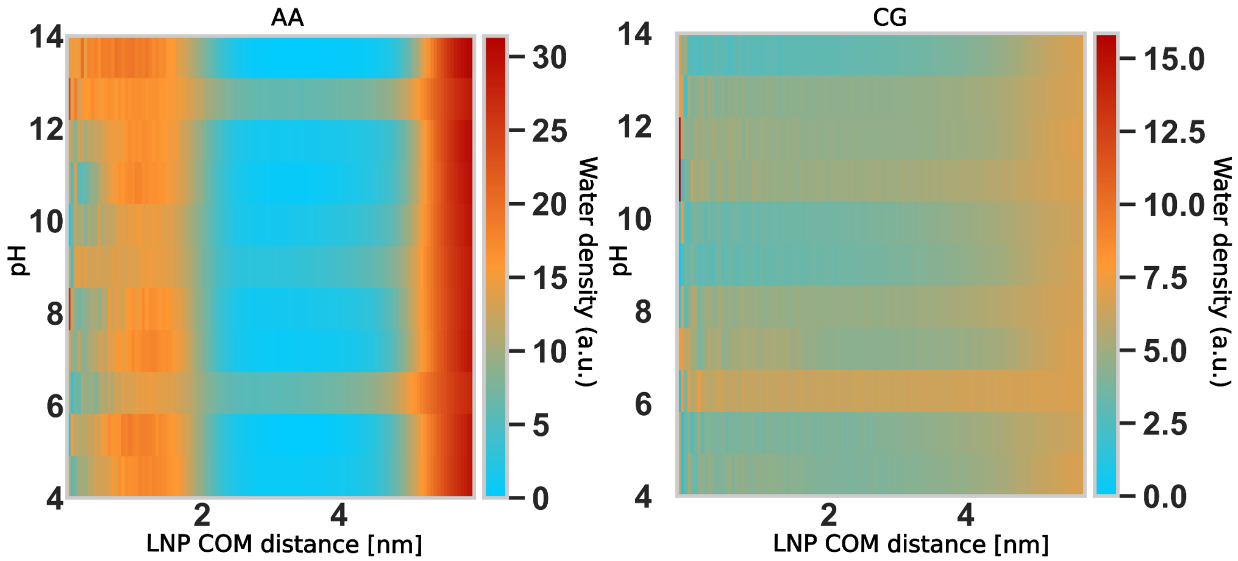


***Fig. S11: Water-density profiles as a function of distance from the LNP center of mass (COM) at different pH values.***

*Water density was computed from the atomistic CpHMD trajectories (left) and from extended coarse-grained (CG) simulations initiated from backmapped atomistic structures (right), and averaged over replicas for each pH. In atomistic, across all conditions, the LNP core—where the siRNA resides—remains well hydrated. In contrast, the outer region shows a slight pH dependence: at acidic pH, the particle remains more hydrated and slightly more open, whereas at higher pH, the water density near the surface slightly decreases, reflecting the more compact and ordered LNP structure observed at elevated pH. The CG results reproduce similar qualitative hydration trends while smoothing short-range fluctuations due to longer timescale sampling.*

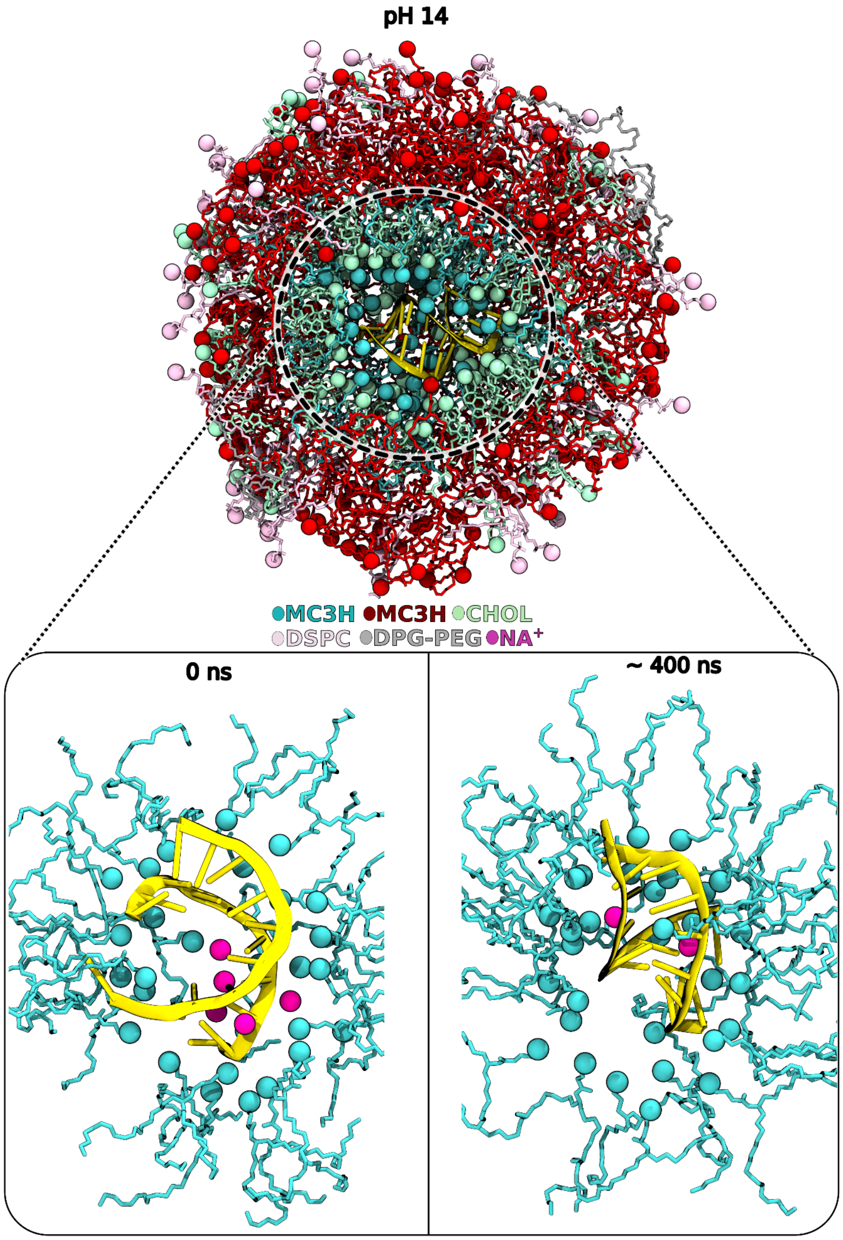


***Fig. S12: Representative CpHMD snapshot at pH 14 illustrating the persistence of protonated MC3 residue near the RNA core.***

*Even at highly alkaline pH (14), the MC3 residues located in the RNA-proximal region remain protonated (MC3H), consistent with the elevated local pKa reported in Fig. 2d. The bottom insets show a zoomed view of the RNA environment at the start of the simulation (t = 0, left) and after ~400 ns (right). Although several $Na^{+}$ ions initially occupy the vicinity of the RNA backbone, they are progressively displaced by MC3H headgroups, which migrate inward and form stronger electrostatic and water-mediated interactions with the phosphate groups. This behavior highlights that monovalent counterions cannot compete with the cationic MC3H for stabilizing the RNA microenvironment, explaining why the inner MC3 shell remains protonated even at extreme pH.*

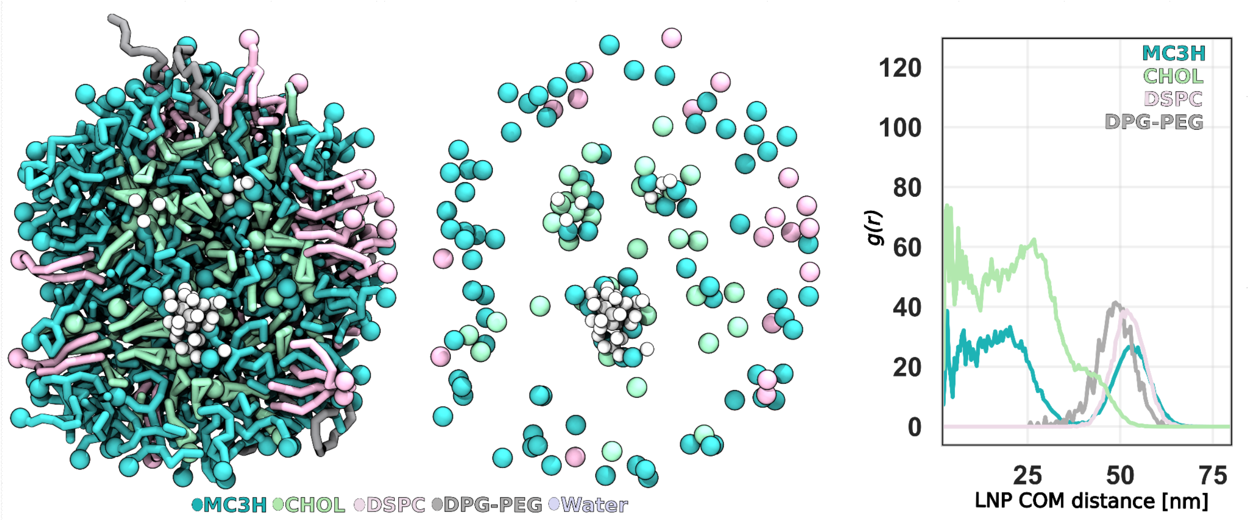


***Fig. S13: Representative structures of the self-assembled empty LNP (no siRNA).***

***Left****: cross-sectional view of the empty LNP from CG self-assembly, showing that in the absence of RNA, the particle adopts a more open, bleb-like internal organization features.* ***Middle****: Visualization of only the lipid headgroups and water highlights the irregular packing and internal water.* ***Right:*** *Radial distribution functions (RDFs) of lipid headgroups around LNP center of mass (COM).*

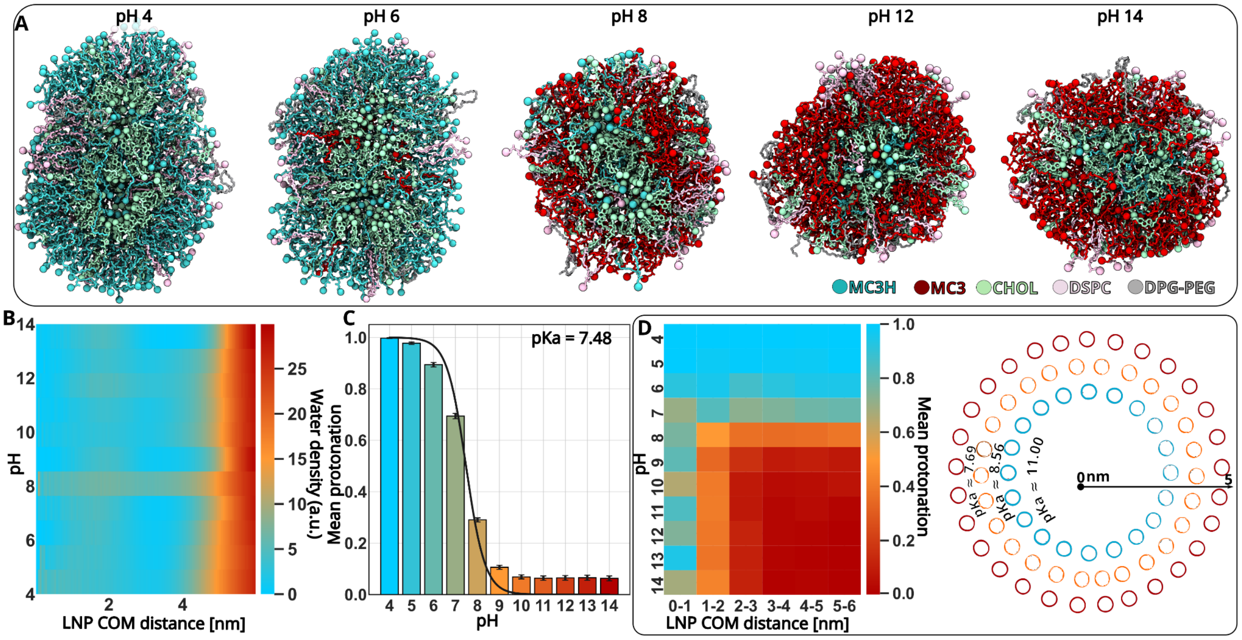


***Fig. S14: Structural and protonation behavior of empty LNPs (without cargo/siRNA).***

*(**A**) Representative snapshots of the atomistic empty LNP at selected pH values from CpHMD simulations. Unlike the RNA-loaded particle, the empty LNP adopts a more open, bleb-like internal arrangement with some showing more than one internal water pocket. Nevertheless, the empty particle also becomes systematically more compact as pH increases (compare low → high pH). (**B**) Distance-resolved water density as a function of LNP center-of-mass (COM) distance and pH. The empty particle shows progressive dehydration at distances beyond ~4 nm as pH rises, consistent with the global compaction seen in the snapshots. (**C**) Global titration of MC3 residues in the empty LNP obtained from the CpHMD (mean protonation fraction ± SD. across two replicas, see Method). Fitting the titration curve to the Henderson–Hasselbalch relation yields an apparent pKa ≈ 7.48 for MC3 in the empty formulation — not significantly different from the value (7.72) found for the siRNA-loaded particle. Convergence of pKa is presented in Fig. S15. (**D**) **Left**: Mean MC3 protonation fraction binned by radial distance from the particle COM and by pH. Although the global titration is similar to the loaded system, the inner region of the empty particle is less protonated than the RNA-stabilized core (compare to Fig. 2). **Right**: converting these distance-dependent titration curves into local apparent pKa values shows a high pKa in the very center (~11) but a rapid drop to ~8.5 just outside the core. Colored circles (red, orange, and blue) indicate the mean protonation state at pHs higher than 9, with higher mean protonation closer to the RNA and decreasing protonation with increasing radial distance.*

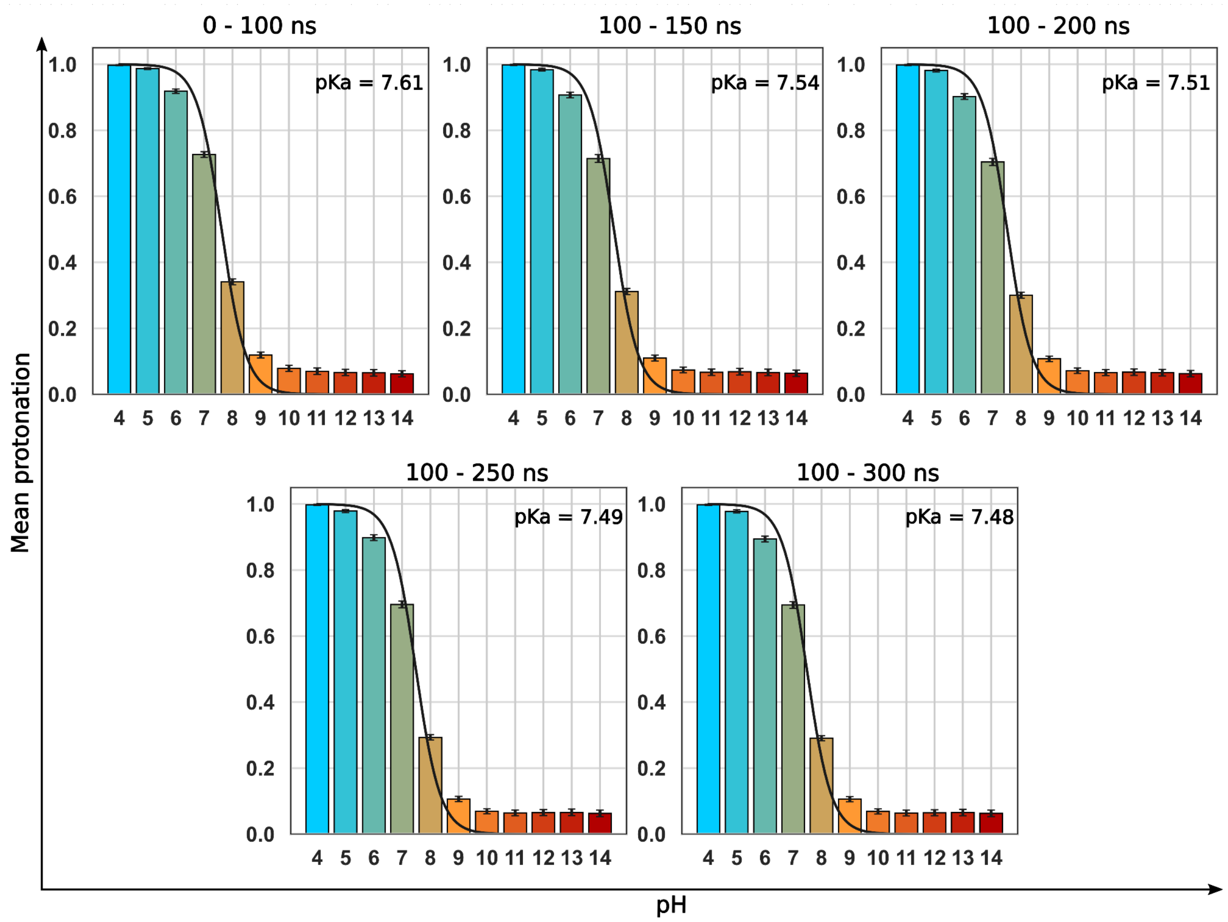


***Fig. S15: Convergence of MC3 protonation behavior in empty LNP.***

*Mean protonation fraction of MC3 residues at each pH value (4–14) shown together with the corresponding titration curve fitted using the Henderson–Hasselbalch equation, calculated over two replicas and at time intervals of 0–100 ns, 100–150 ns, 100–200 ns, 100–250 ns, and 100–300 ns—used to assess convergence.*

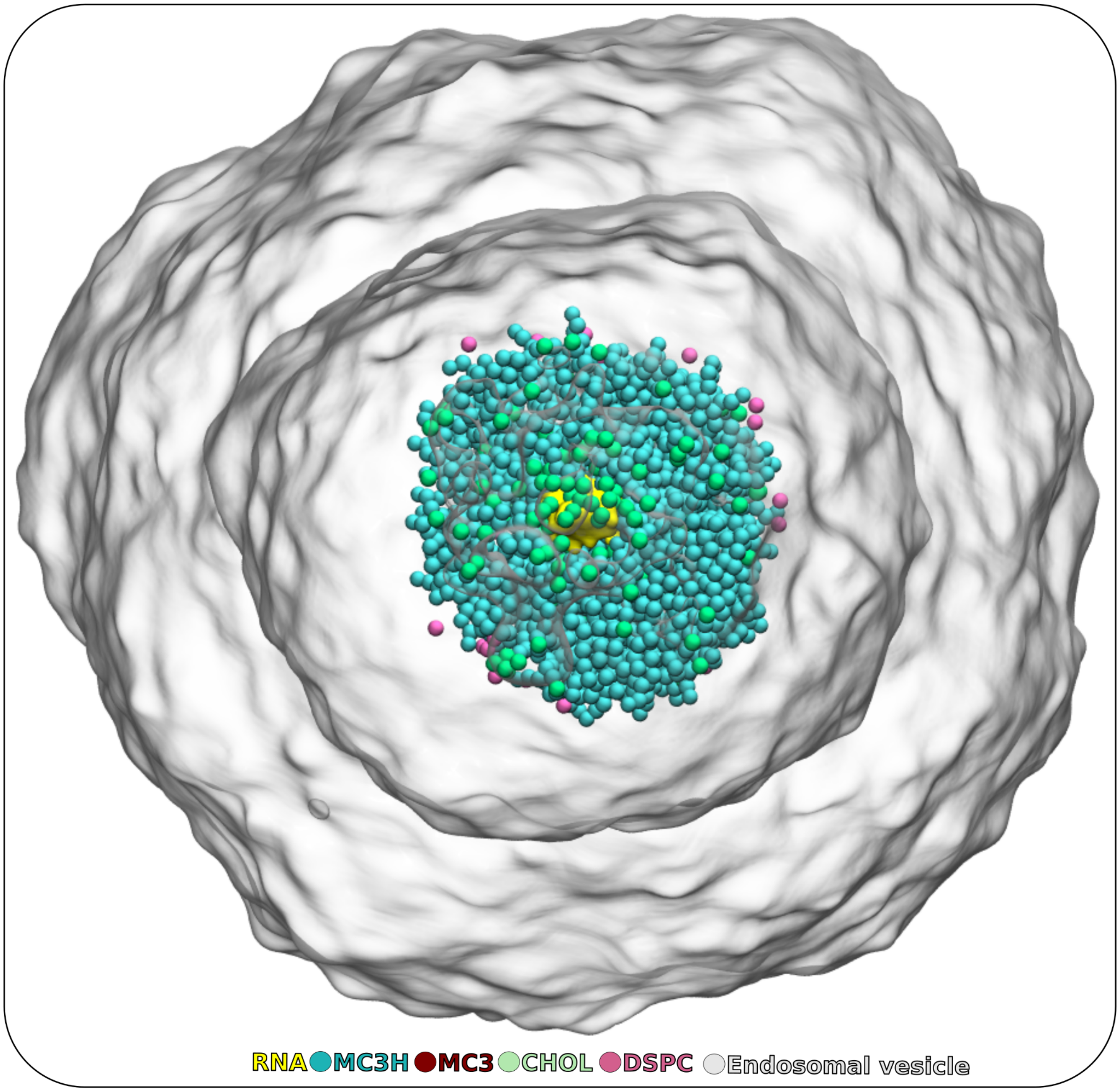


***Fig. S16: Initial configuration of the LNP inside the model endosomal vesicle.***

*A representative starting snapshot showing the full endosomal vesicle (rendered as a smooth silver surface) with the LNP placed in the lumen prior to CG simulation. The LNP components are colored as: siRNA (gold), protonated MC3H (cyan), neutral MC3 (red), cholesterol (mint), DSPC (pink), and DPG-PEG (gray).*

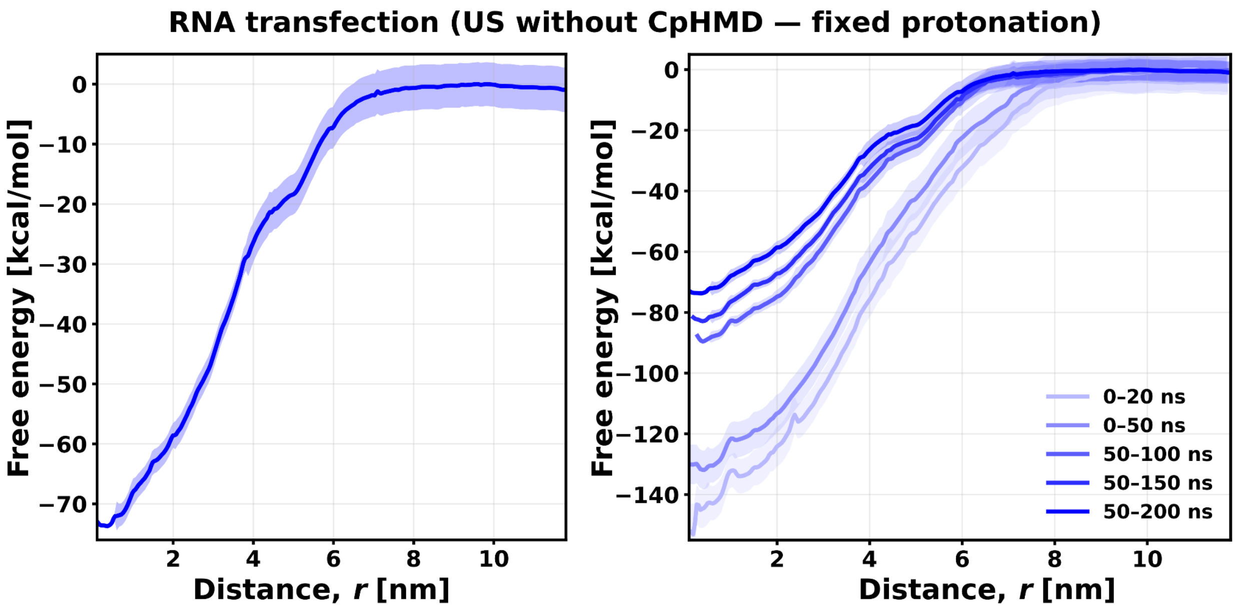


***Fig. S17: Free-energy profile for siRNA translocation using fixed-protonation umbrella sampling (without CpHMD), and its convergence behavior.***

*(**Left**) Potential of mean force (PMF) obtained from umbrella sampling using a fixed protonation state for all MC3 lipids. The profile shows a huge barrier (~72 kcal/mol), indicating that treating MC3 protonation as static is not suitable for this process, as the strong, dynamically evolving electrostatic interactions between MC3 and the RNA dominate the energetics. Note that the time convergence profile (**right**) suggest that this cannot be fully explained as a simple sampling problem, suggesting that strong electrostatic interactions implicit to the use of a fixed charge state for M3C leads to these unrealistic results.*

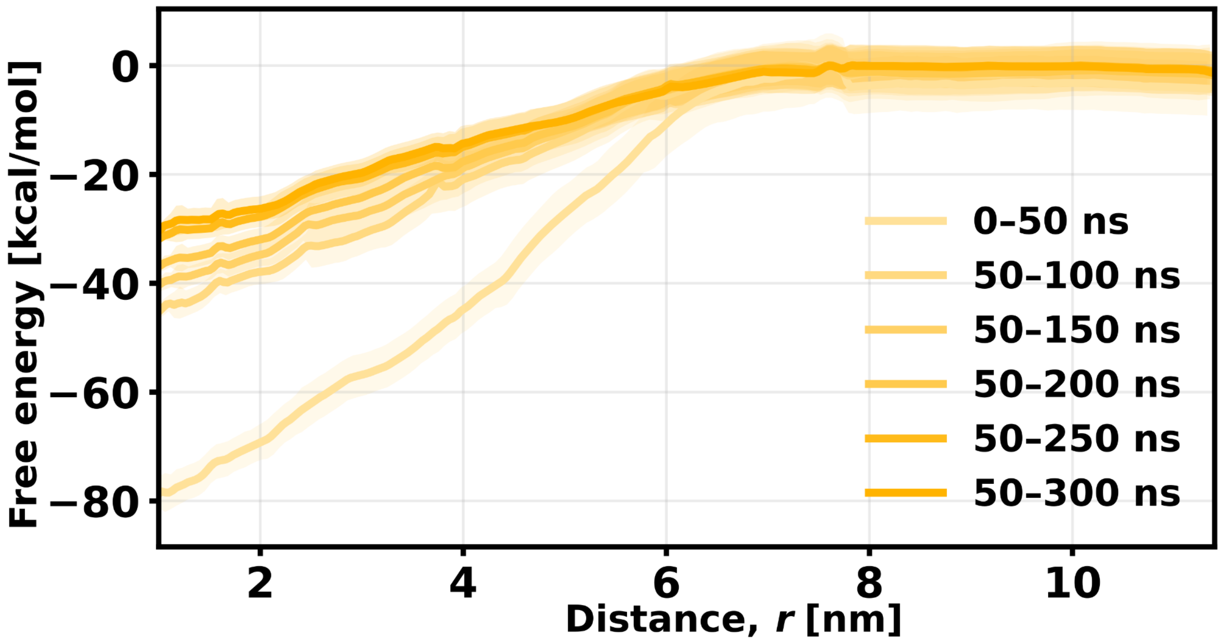


***Fig. S18: Free-energy convergence for siRNA translocation using dynamic protonation of MC3 lipids (CpHMD-US)***.

*Potential of mean force (PMF) obtained from umbrella sampling using dynamic protonation state for all MC3 lipids, i.e., coupled with constant-pH MD (CpHMD).*

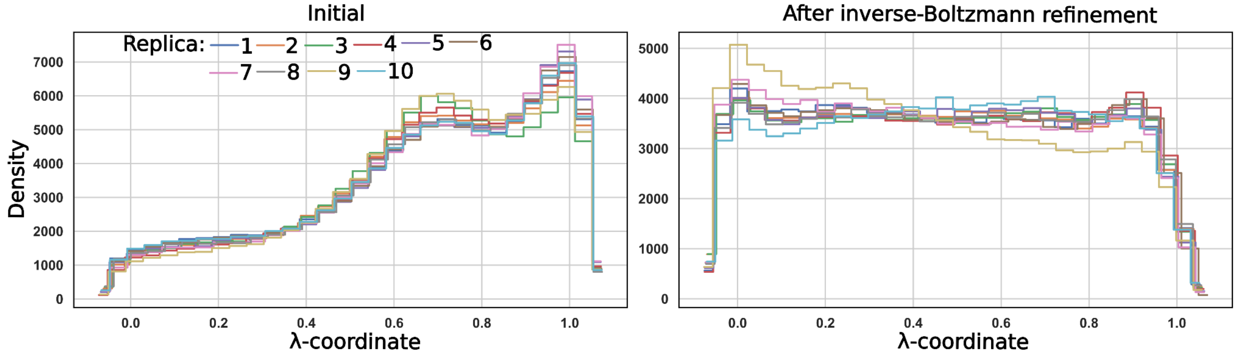


***Fig. S19: Calibration of MC3 parameters for CpHMD.***

*(**Left**) The initial estimate of the MC3 dV/dλ correction term gives λ-distributions that are reproducible across replicates but not yet flat at pH = pKa, indicating that the protonation coordinate is still biased.*

*(**Right**) After applying inverse-Boltzmann refinement and rerunning the calibration simulations, the λ-distributions become both consistent and essentially flat, confirming that the final MC3 parameter set satisfies the expected CpHMD conditions and is suitable for production simulations.*
